# Dynamics of ribosomes and release factors during translation termination in *E. coli*

**DOI:** 10.1101/243485

**Authors:** Sarah Adio, Heena Sharma, Tamara Senyushkina, Prajwal Karki, Cristina Maracci, Ingo Wohlgemuth, Wolf Holtkamp, Frank Peske, Marina V. Rodnina

## Abstract

Release factors RF1 and RF2 promote hydrolysis of peptidyl-tRNA during translation termination. The GTPase RF3 promotes recycling of RF1 and RF2. Using single molecule FRET together with ensemble kinetics, we show that ribosome termination complexes that carry two factors, RF1–RF3 or RF2–RF3, are dynamic and fluctuate between non-rotated and rotated states, while each factor alone has its distinct signature on the ribosome dynamics and conformation. Dissociation of RF1 depends on peptide release and the presence of RF3, whereas RF2 can dissociate spontaneously. RF3 binds in the GTP-bound state and can rapidly dissociate without GTP hydrolysis from termination complex carrying RF1. GTP cleavage helps RF3 release from ribosomes stalled in the rotated state in the absence of RF1. Our data suggest how the stochastic assembly of the ribosome–RF1–RF3–GTP complex, peptide release, and ribosome fluctuations promote termination of protein synthesis and recycling of the release factors.

## Introduction

Termination of protein synthesis occurs when a translating ribosome encounters one of the three universally conserved stop codons UAA, UAG or UGA. In bacteria the release of the nascent peptide is promoted by release factors RF1 and RF2 which recognize the stop codons in the A site and hydrolyze the ester bond in the peptidyl-tRNA bound to the P site, allowing the nascent peptide to leave the ribosome through the polypeptide exit tunnel (Dunkle and Cate, 2010; Nakamura et al., 1996). RF1 and RF2 bind to the ribosome in the space between the small and large ribosomal subunits. RF1 and RF2 differ in their stop codon specificity: RF1 utilizes a conserved PET motif to recognize UAG and UAA codons, whereas RF2 uses an SPF motif to recognize UGA and UAA codons. Both RF1 and RF2 have a universally conserved GGQ motif which promotes the catalysis of peptidyl-tRNA hydrolysis (Seit-Nebi et al., 2001); mutations of the GGQ motif to GAQ or GGA inhibit peptide release (Frolova et al., 1999; Mora et al., 2003; Shaw and Green, 2007; Zavialov et al., 2002). After peptide release, RF1 and RF2 dissociate from the post-termination complex to allow for the next steps of translation. The dissociation is accelerated by RF3, a translational GTPase that binds and hydrolyses GTP in the course of termination (Freistroffer et al., 1997; Koutmou et al., 2014; Zavialov et al., 2002).

The mechanism of termination is unclear and there are two different models concerning the sequence of events, including the timing of peptide release, the order of RF1, RF2 and RF3 binding and dissociation, and the role of nucleotide exchange in RF3 and GTP hydrolysis. The first model of translation termination, which was proposed by Ehrenberg and colleagues (Zavialov et al., 2001; Zavialov et al., 2002), is based on the following key observations. The authors reported that free RF3 has a much higher affinity for GDP (K_d_=5.5 nM) than for GTP (K_d_=2.5 μM) or GDPNP (K_d_=8.5 μM) (Zavialov et al., 2001), which would imply that at cellular GTP/GDP concentrations RF3 is expected to be predominantly in the GDP form. The exchange of GDP for GTP occurs only when RF3–GDP binds to the ribosome in complex with RF1 or RF2 (Zavialov et al., 2001). In the absence of the nucleotide, RF3–dependent RF1/2 recycling is slow, which has been interpreted as an indication for a high affinity complex of apo-RF3 to the ribosome–RF1/2 complex (Zavialov et al., 2001). Furthermore, because RF3–dependent turnover GTPase activity was stimulated by peptidyl-tRNA hydrolysis, the authors suggested that RF3 binds to the ribosome–RF1 complex only after the peptide is released. Based on these results, Ehrenberg et al. suggested the following sequential mechanism of termination: RF1/RF2 bind to the ribosome and hydrolyze peptidyl-tRNA, allowing RF3–GDP to enter the ribosome occupied by RF1 or RF2 to form an unstable encounter complex. Dissociation of GDP leads to a stable high affinity complex with RF3 in the nucleotide-free state. The subsequent binding of GTP by RF3 promotes RF1/RF2 dissociation. In the final step, RF3 hydrolyses GTP and as a result dissociates from the ribosome (Zavialov et al., 2001; Zavialov et al., 2002).

An alternative model was proposed when it turned out that the affinity of RF3 to GDP and GTP is on the same order of magnitude (5 nM and 20 nM, respectively (Koutmou et al., 2014; Peske et al., 2014)). As the cellular GTP concentration is at least ten times higher than the GDP concentration (Bennett et al., 2009), these affinities imply that nucleotide exchange in RF3 can occur spontaneously, off the ribosome, and thus RF3 could enter the ribosome in either the GTP- or GDP-bound form. Consistent with previous findings (Zavialov et al., 2001; Zavialov et al., 2002), ribosome–RF1 or ribosome–RF2 complexes accelerate nucleotide exchange in RF3 (Koutmou et al., 2014; Peske et al., 2014); however, this effect is independent of peptide release, because also a catalytically inactive RF2 mutant activates nucleotide exchange in RF3 (Peske et al., 2014; Zavialov et al., 2002). Binding of GTP to RF3 in the complex with the ribosome and RF2 is rapid (130 s^-1^) (Peske et al., 2014), and thus the lifetime of the apo-RF3 state would be too short to assume a tentative physiological role. Peptide release results in the stabilization of the RF3–GTP–ribosome complex, thereby promoting the dissociation of RF1/2, followed by GTP hydrolysis and dissociation of RF3–GDP from the ribosome (Peske et al., 2014).

Efficient translation termination requires not only the coordinated action of the release factors, but also entails conformational dynamics of the factors and the ribosome. The key conformational motions of the ribosome identified so far include the rotation of ribosomal subunits relative to each other, the swiveling motion of the body and head domains of the small ribosomal subunit, and the movement of the ribosomal protein L1 in and out relative to the E-site tRNA. During the elongation cycle these motions are gated by ligands such as translation factors and tRNAs (Adio et al., 2015; Chen et al., 2011; Cornish et al., 2008; Horan and Noller, 2007; Sharma et al., 2016; Shi and Joseph, 2016; Valle et al., 2003; Wasserman et al., 2016). Crystal structures show that termination complexes with RF1 or RF2 are predominantly in the non-rotated (N) state irrespective of peptide release, that is in pre- and post-hydrolysis complexes (PreHC and PostHC, respectively); the P-site tRNA in the complexes is in the classical state and the L1 stalk in an open conformation (Jin et al., 2010; Korostelev et al., 2008; Laurberg et al., 2008; Weixlbaumer et al., 2008). Also a single molecule fluorescence resonance energy transfer (smFRET) study using donor and acceptor FRET pairs in the tRNA, RF1 and L1 showed that binding of RF1 to termination complexes results in the stabilization of a state with the tRNA in a classical state and an open conformation of the L1 stalk (Sternberg et al., 2009); the rotation of the ribosomal subunits was not investigated directly in that study. The high sequence similarity between RF1 and RF2 suggests that the two factors interact with the ribosome in the same manner and promote peptide release by a similar mechanism (Freistroffer et al., 1997; Zavialov et al., 2001). However, structures of RF1 or RF2 bound to PostHC reveal differences regarding the interaction with the L11 region of the 50S subunit (Korostelev et al., 2008; Laurberg et al., 2008; Petry et al., 2005; Pierson et al., 2016; Rawat et al., 2006; Rawat et al., 2003; Weixlbaumer et al., 2008). Thus, it is not clear whether RF1 and RF2 follow the same mechanism and whether they respond in the same way to the recruitment of RF3 to termination complexes.

Binding of RF3 with a non-hydrolyzable GTP analog, GDPNP, to PostHC in the absence of RF1/RF2 induces the formation of the rotated (R) state of the ribosome with the P-site tRNA in a hybrid state and the closed conformation of the L1 stalk (Gao et al., 2007; Jin et al., 2011; Zhou et al., 2012). A very similar effect of RF3–GDPNP on subunit rotation was found by smFRET (Sternberg et al., 2009). Modeling of the atomic structures of RF1 and RF2 into the cryoEM structure of RF3–bound PostHC suggests that the RF3–induced ribosome rearrangements break the interactions of RF1/RF2 with both the decoding center and the L11 region of the ribosome, leading to the release of the RF1/RF2 (Gao et al., 2007). However, in the PostHC with RF1 and RF3 in the apo form, i.e., in the absence of added nucleotide, the conformation of the ribosome represents the N state, whereas the position of RF1 changes in response to RF3 binding (Pallesen et al., 2013). smFRET measurements carried out with a PostHC in the presence of excess RF1 showed that RF3–GTP induced short-lived excursions into the L1-closed (presumably R) state, but generally the L1-open (and likely N) state was stabilized (Sternberg et al., 2009). The interaction of RF3 with the PreHC has not been studied.

Here we use five different FRET pairs to monitor subunit rotation, the orientation of the P-site tRNA–L1 elements, and the interactions of RF1, RF2, and RF3 with the ribosome during termination by single molecule TIRF microscopy. Our results demonstrate how the recruitment of release factors change the ribosome conformation in the PreHC and PostHC, how the dissociation of the factors is achieved, show the differences in the function of RF1 and RF2, and explain the importance of GTP binding and hydrolysis.

## Results

### RF1 and RF2 have distinct effects on ribosome dynamics

To monitor the rotation of the ribosomal subunits with respect to one another, we utilized fluorescent labels attached to the proteins S6 and L9, S6-Cy5 and L9-Cy3, respectively. This FRET pair has been extensively characterized in both smFRET and ensemble kinetics experiments and senses the formation of the N or R state of the ribosome (Cornish et al., 2008; Ermolenko et al., 2013; Sharma et al., 2016). We prepared termination complexes on an mRNA which is translated up to the stop codon UAA recognized by both RF1 and RF2. The complexes contain a peptidyl-tRNA in the P site and have a stop codon in the A site; those complexes are denoted as PreHC. In the absence of termination factors, PreHC is found predominantly in the high FRET state (0.7) corresponding to the N state (**Figures 1A, Figure 1-Figure Supplement 1; Table S1**), consistent with previous reports (Cornish et al., 2008; Qin et al., 2014; Sharma et al., 2016). The traces are static, i.e., we do not observe transitions between N and R states within the limits given by the time resolution of the camera and the finite experimental observation time. Distribution of FRET states and conformational dynamics of PreHC are independent of the tRNA in the P site and of the presence of a single N-terminal amino acid (fMet) or a dipeptide (fMetPhe, fMetVal or fMetLys) at the P-site-tRNA (**Figure 1-Figure Supplement 1; Table S1**). This allowed us to use the PreHC with fMet-tRNA^fMet^ in the P site and a stop codon in the A site as a minimal model system, following previous publications that suggested that fMet-Stop is a good model system to study the mechanism of translation termination (Koutmou et al., 2014; Kuhlenkoetter et al., 2011; Pierson et al., 2016; Shi and Joseph, 2016; Sternberg et al., 2009).

**Figure 1.**
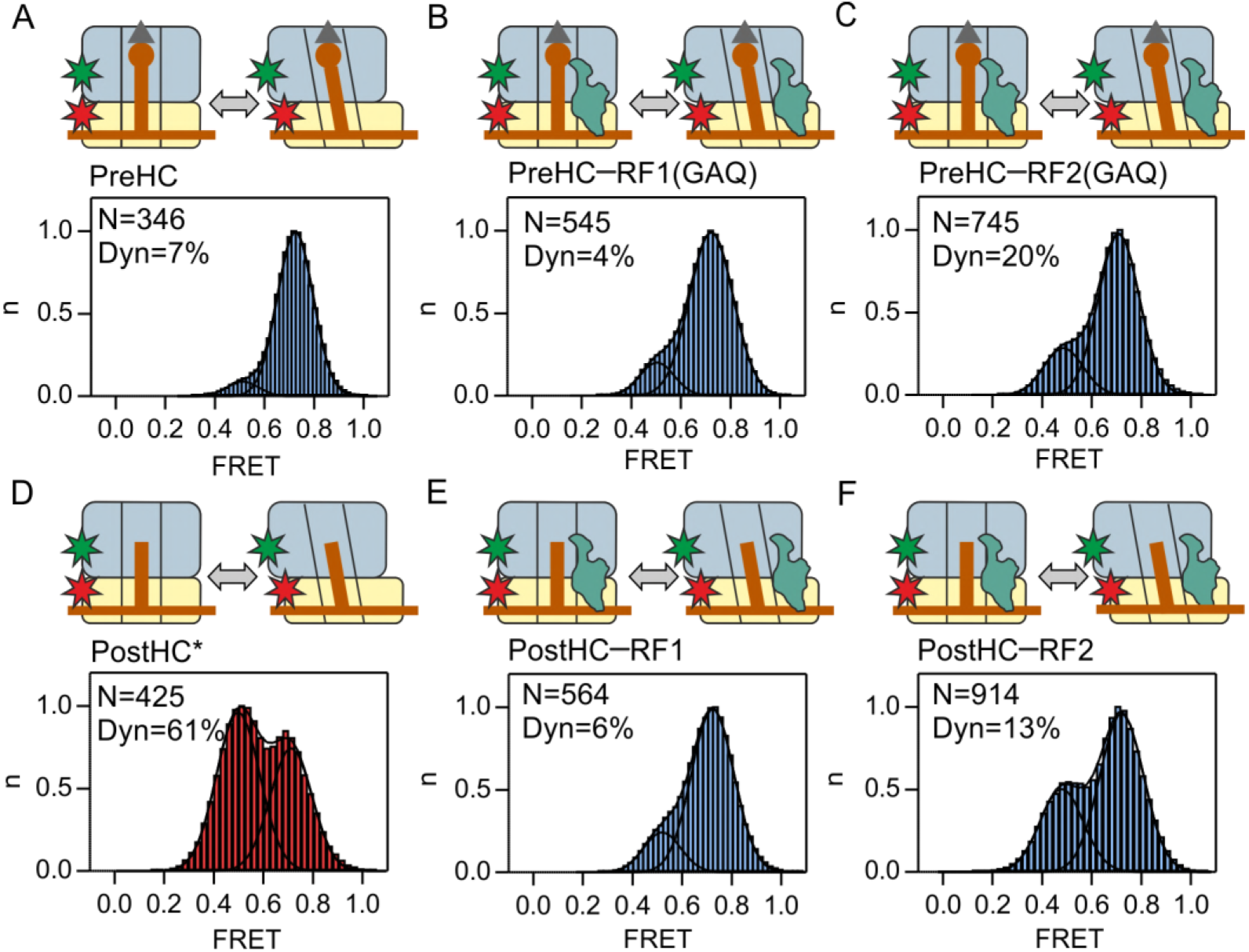
Subunit rotation monitored using FRET between S6-Cy5 and L9-Cy3. (A,D) Rotation in the absence of release factors. (B,E) In the presence of RF1 or RF1(GAQ) (1 μM). (C,F) In the presence of RF2 or RF2(GAQ) (1 μM). Cartoons show the complex composition. Grey triangles and brown circles represent the formyl group and the amino acid of fMet, respectively; stars indicate the positions of the Cy3 (green) and Cy5 (red) labels. PostHC* was generated by addition of puromycin to PreHC. Histograms display the normalized distribution of FRET states with Gaussian fits showing the distribution between N (FRET 0.7) and R (FRET 0.5) states. N is the number of traces entering the histogram. Dyn is the percentage of dynamic complexes that show transitions between N and R states. Static complexes (with 0–20% traces showing fluctuations) are represented by blue histograms, moderately dynamic complexes (20–45%) are shown in pink, highly dynamic complexes (>45%) are shown in dark red. *See also Figure 1-Figure Supplement 1, Figure 1-Figure Supplement 2 Figure 1-Figure Supplement 3 and Table S1*

To study the effect of RF1/RF2 binding on subunit rotation, we added an excess of the factors sufficient to saturate the ribosomes (Zavialov et al., 2002). Peptide release was avoided by using RF1(GAQ) or RF2(GAQ) mutants which are catalytically deficient (Frolova et al., 1999; Zavialov et al., 2002) (**Figure 1-Figure Supplement 2A**). Binding of RF1(GAQ) to PreHC emphasizes the N state (**Figure 1B**). Surprisingly, RF2(GAQ) binding to PreHC results in a somewhat larger portion of dynamic ribosomes and of R state (**Figure 1C**). The ability of peptidyl-tRNA to adopt R/hybrid state at room temperature has been demonstrated previously by cryo-EM (Fischer et al., 2010)

To prepare PostHC without using termination factors, we released nascent peptides with the help of puromycin, an analog of the A-site aminoacyl-tRNA that reacts with the peptidyl-tRNA in the P site to form peptidyl-puromycin (which then dissociates from the ribosome) and a deacylated tRNA in the P site. FRET time courses of PostHC in the absence of the factors show transitions between the high 0.7 and low 0.5 FRET indicating a dynamic equilibrium between N and R states (**Figure 1D**). The exact distribution of states depends on the P-site tRNA (**Figure 1-Figure Supplement 1; Table S1**) (Fei et al., 2011), with the fMet-stop complex behaving similarly to that with fMetVal-tRNA^Val^, thus underscoring the suitability of the minimal model system. RF1 halts the complex in the N state (**Figure 1E**, **Figure 1-Figure Supplement 2B,D**), in agreement with the previous smFRET study suggesting that RF1 binding stabilizes the L1 stalk in the open conformation characteristic for the N state (Sternberg et al., 2009). Complexes with RF1 are static, i.e., there are very few transitions between N and R states. Binding of RF2 to PostHC shifts the equilibrium towards the N state, but not to the same extent as binding of RF1 (**Figure 1F**, **Figure 1-Figure Supplement 2C,E**). Complexes with RF2 make occasional N to R transitions, in particular with RF2(GAQ) bound to PostHC (**Figure1-Figure Supplement 2C**).

To further probe the differences between RF1 and RF2, we performed the experiment in the presence of limiting amounts of factors and monitored ribosome rotation in real time. RF1 binding to PreHC does not change the FRET efficiency appreciably, as the complex is predominantly in the N state with or without the factor (**Figure 1-Figure Supplement 3A,B; Table S1**). Stabilization of the N state by RF1 is independent of peptide release, because the distribution of the N and R state is identical with wild-type RF1 and the RF1(GAQ) mutant. Binding of RF1 to PostHC halts fluctuating ribosomes in the N state (**Figure 1-Figure Supplement 3C**). Synchronization of time courses to the last R to N transition indicates that RF1 binding is rapid and occurs within the acquisition time of one camera frame; the binding event of RF1 to the N state is not visible, because it does not result in a FRET change.

In contrast to RF1, RF2 binding to PreHC and peptide release appears to shift the N to R equilibrium towards the R state (**Figure 1-Figure Supplement 3D,F**). The resulting PostHC with RF2 is dynamic and fluctuates between N and R states as shown by synchronization of FRET traces to the first N to R transition. These transitions depend on peptide release, as the interaction of RF2(GAQ) with PreHC does not result in a signal change (**Figure 1-Figure Supplement 3E**). These data suggest that RF1 and RF2 have distinct effects on the ribosome dynamics and, possibly, different dwell times on the ribosome.

### Retention of RF1 and RF2 on the ribosome

To measure how long the factors remain bound to the ribosome, we prepared Cy5-labeled RF1 and RF2, as well as the respective RF1/2(GAQ) mutants and ribosomes containing Cy3-labeled protein L11 (Adio et al., 2015; Chen et al., 2011; Geggier et al., 2010; Holmberg and Noller, 1999; Stoffler et al., 1980) (**Figure 2-Figure Supplement 1A**). The biochemical activity of labeled release factors was indistinguishable from that of the unlabeled counterparts (**Figure 2-Figure Supplement 1B,C**) and the factors were fully methylated (**Figure 2-Figure Supplement 2**). L11 constitutes part of the factor binding site (Pallesen et al., 2013; Peske et al., 2014; Petry et al., 2005). Recruitment of the factors to the ribosome is expected to result in efficient FRET between the donor and acceptor dyes. Binding of RF1 or RF1(GAQ) to either PreHC or PostHC results in a single population of ribosome complexes with a mean FRET efficiency of 0.7 (**Figure 2A-C**). RF1 or RF1(GAQ) remain stably bound to the ribosome independent of peptide release. The duration of FRET signals is limited only by the photobleaching rate at the given imaging condition and the estimated upper limit of the dissociation rate constant is 0.2 s^-1^ (**Table S1**). Binding of RF2 or RF2(GAQ) to PreHC and PostHC also leads to a FRET efficiency of about 0.7 (**Figure 2D-F**). However, the residence time of RF2 is much shorter compared to RF1 or RF1(GAQ), with the k_off_ values in the range of 0.8–1.3 s^-1^ (**Figure 2D-F**, **Table S1**).

**Figure 2.**
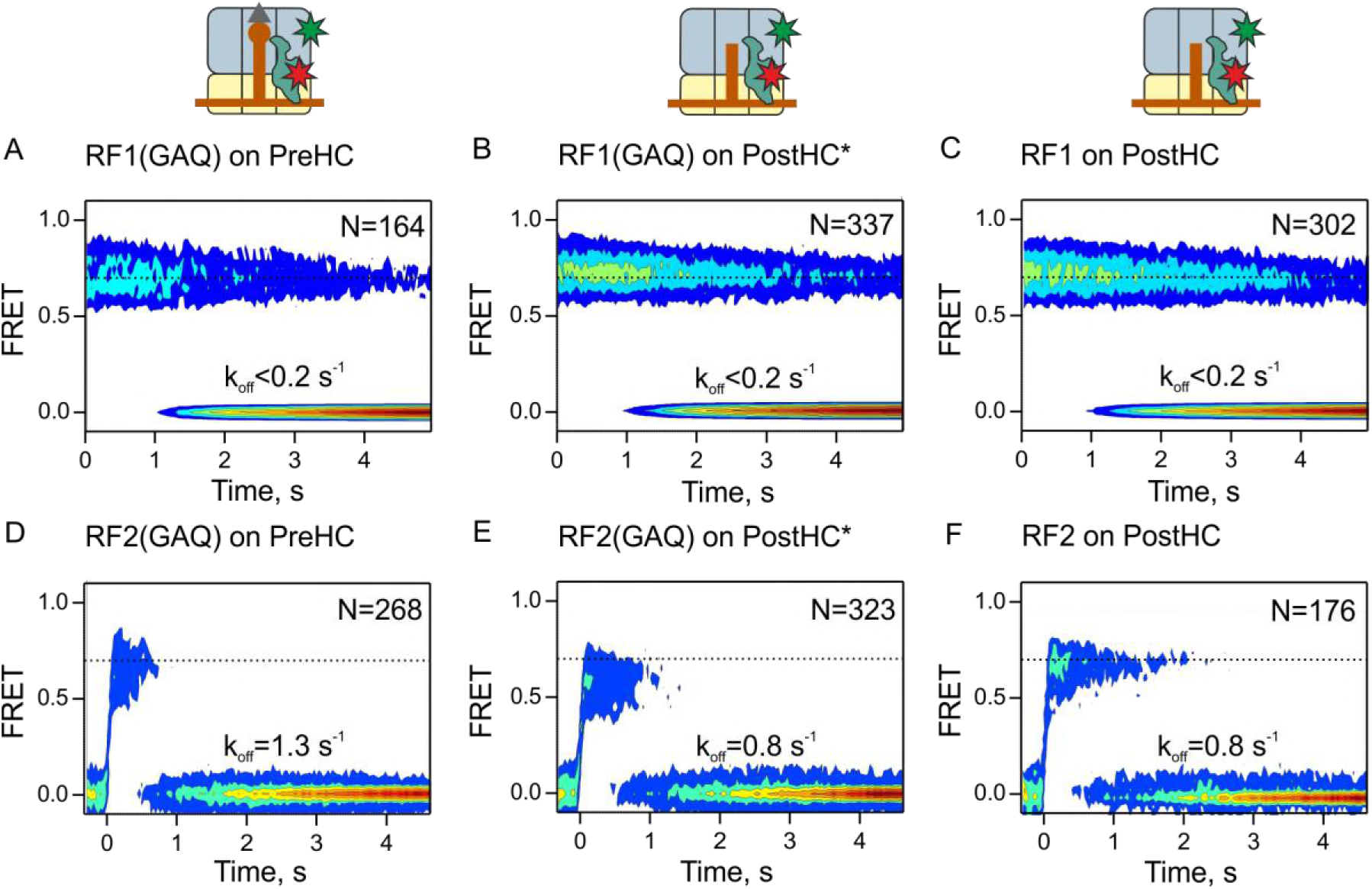
Residence times of RF1 and RF2 on PreHC and PostHC. (A-C) smFRET upon addition of RF1-Cy5 or RF1(GAQ)-Cy5 to PreHC or PostHC labeled at protein L11 with Cy3. (D-F) smFRET upon addition of RF2-Cy5 or RF2(GAQ)-Cy5 to PreHC or PostHC labeled at protein L11 with Cy3. Experiments were carried out with catalytic amounts of labeled release factors (10 nM). Individual traces were combined in contour plots and for RF2 synchronized to the first FRET event. N indicates the number of individual traces in the contour plot. k_0_ff is the rate of RF1 or RF2 dissociation. *See also Figure 2-Figure Supplement 1, Figure 2-Figure Supplement 2, Figure 2-Figure Supplement 3 and Table S1*

The difference in the dissociation rates of RF1 and RF2 suggests that RF1 needs an auxiliary factor, RF3, to help it to dissociate from the ribosome, whereas RF2 recycles independently of RF3. This notion is consistent with previous reports (Petropoulos et al., 2014; Zavialov et al., 2002) and is further supported by our peptide hydrolysis turnover assay (**Figure 2-Figure Supplement 3E**). When RF1 is added in catalytic amounts, very little peptide is released in the absence of RF3, i.e., when RF1 turnover depends on its intrinsic dissociation rate from the ribosome. In the presence of RF3 peptide release is very efficient. In contrast, catalytic amounts of RF2 are sufficient to release peptide even in the absence of RF3, although addition of RF3 facilitates the reaction. Addition of RF3 to the PreHC–RF2 increases the fraction of the R states compared to the complex with RF2 alone and enhances fluctuations between the states (**Figure 2-Figure Supplement 3A,C**). Most of the dynamic ribosomes fluctuate rapidly. After peptide release the preference for the R state does not change, but the number of dynamic ribosomes decreases and the transitions are slower (**Figure 2-Figure Supplement 3B,D; Table S1**). Thus, RF3 is essential for RF1, but not for RF2 recycling.

### Interaction with RF3–GTP

RF3 has the opposite effect on subunit rotation than RF1 or RF2 (**Figure 3**). Binding of RF3 to PreHC, which in the absence of the factor is predominantly static and favors the N state, strongly shifts the equilibrium towards the R state (**Figure 3A**; blue line shows the distribution of states in the absence of RF3). About 50% of the traces are now dynamic and show reversible N to R transitions. The dwell time distributions of both N and R states are biphasic: The majority (~80%) shows rapid N to R and R to N transitions (k_N→R_ = 6.2 s^-1^, k_R→N_ = 2.2 s^-1^), whereas about 20% show slower N to R (1.5 s^-1^) and R to N (0.7 s^-1^) transitions (**Table S1**). This finding seems unexpected, given that peptidyl-tRNA is expected to favor the N/classical state. However, previous cryo-EM reconstructions indicated that peptidyl-tRNA can, in fact, adopt the R state at elevated temperatures (Fischer et al., 2010). At 18°C, close to 22°C used in our experiments, the distribution of the rotation states for the ribosome complex with peptidyl-tRNA (fMetVal-tRNA^Val^) was bimodal, with comparable peaks of N and R states (Fischer et al., 2010). Thus, RF3 appears to bias spontaneous fluctuations of the tRNA in the PreHC, rather than induce a previously disallowed conformation.

**Figure 3.**
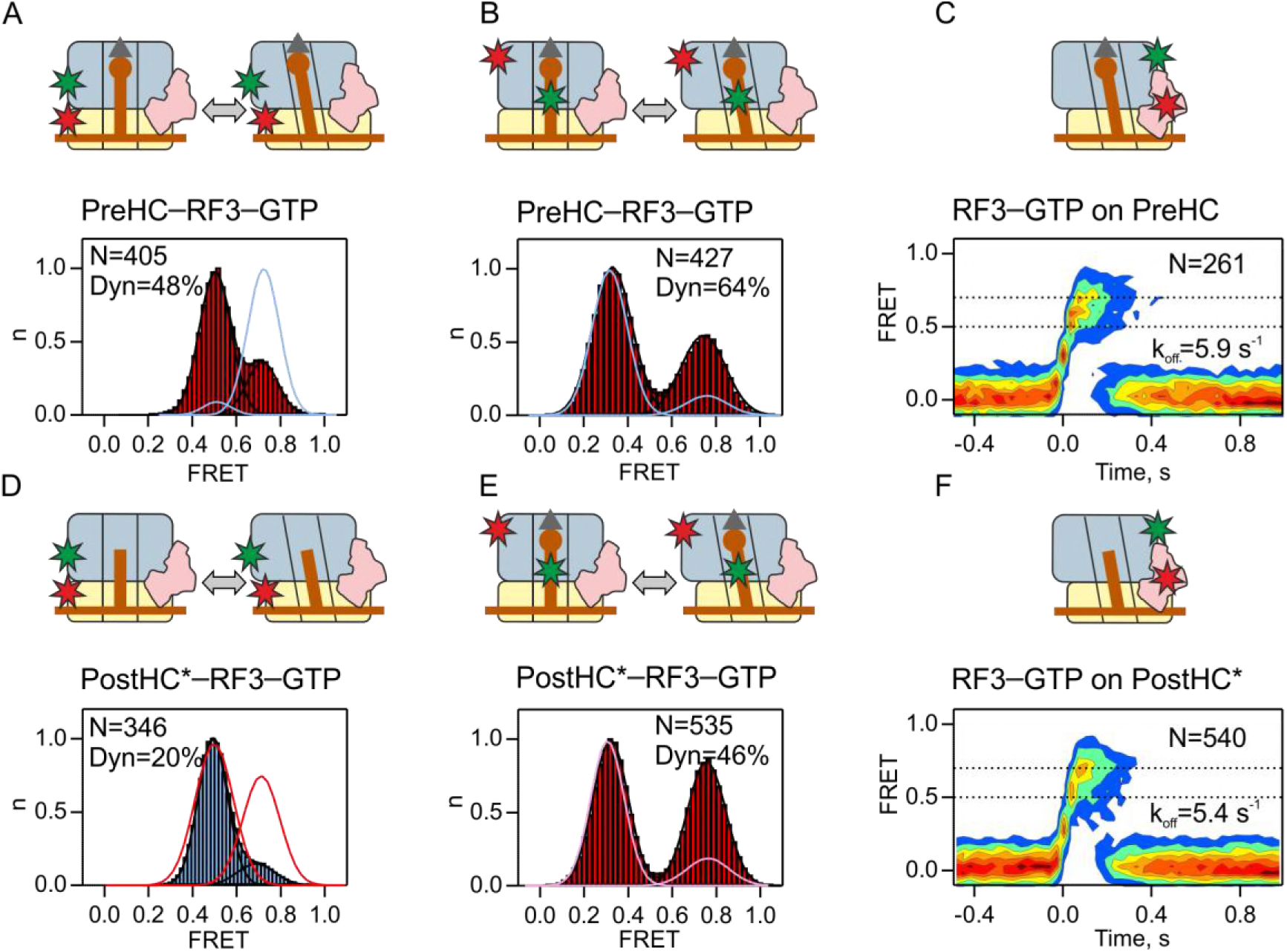
Interaction of RF3–GTP with ribosome complexes. (A,D) Ribosome rotation of PreHC and PostHC* in the presence of excess RF3 (1 μM) with GTP (1 mM). Individual traces (N indicates the number of traces) are combined in a FRET distribution histogram; the percentage of dynamic complexes is indicated (Dyn). Blue line in (A) represents the distribution of FRET states for PreHC in the absence of RF3. Red line in (D) represents the distribution of FRET states for PostHC in the absence of RF3. (B,E) L1–tRNA FRET distribution in PreHC and PostHC* in the presence of excess RF3 (1 μM) with GTP (1 mM). Blue line in (B) represents the FRET distribution for PreHC in the absence of RF3. Pink line in (E) represents the FRET distribution for PostHC in the absence of RF3. (C,F) Contour plots representing the residence time of RF3–Cy5 (10 nM) on PreHC–Cy3 or PostHC-Cy3. Time courses were synchronized to the first FRET event and combined in contour plots. N is the number of traces entering the contour plot. k_off_ is the rate of RF3 dissociation. *See also Figure3-Figure Supplement 1 and Table S1*

To further characterize the conformational changes induced by RF3, we probed the conformation of the P-site tRNA relative to the protein L1. We used the validated FRET pair with the donor label at the tRNA (fMet-tRNA^fMet^(Cy3)) and the acceptor label on the ribosomal protein L1 (L1(Cy5)). The two labels are close together and give a high FRET signal when the L1 stalk is in the closed conformation with the tRNA in the P/E hybrid state and move apart to give low FRET signal when the L1 stalk adopts open conformation with the tRNA in the classic P/P state (Fei et al., 2009; Fei et al., 2008; Munro et al., 2010a; Munro et al., 2010b; Munro et al., 2010c; Sternberg et al., 2009). FRET histograms of PreHC in the absence of RF3 (without RF1/RF2 or in the presence of RF2(GAQ)) are dominated by the low (0.3) FRET population and the traces are static (**Figure 3B**, blue line **Figure 3-Figure Supplement 1A,B** and **Table S1**). This indicates that fMet-tRNA^fMet^ is predominantly in the classic P/P state, in agreement with previous studies (Fischer et al., 2016; Loveland et al., 2017). RF3 induces dynamic transitions between the low (0.3) and high (0.8) FRET state and shifts the equilibrium towards the P/E hybrid state (**Figure 3B**). The P/E hybrid state is short lived and decays rapidly (k_off_ = 6.0 s^-1^**; Table S1**). The decay rate is faster than subunit rotation (kR N = 2.2 s^-1^) suggesting that the two processes are not tightly coupled, consistent with previous notions based on cryo-EM reconstructions (Fischer et al., 2010) and smFRET work (Munro et al., 2010b).

PostHC fluctuates between N/classical and R/hybrid states in the absence of RF3 (**Figure 3D**, red line, (**Figure 3-Figure Supplement 1C**; **Figure 1D**). RF3 halts these fluctuations in the R state (**Figure 3D**), but the transitions between the classic and hybrid states are still observed (**Figure 3E; Table S1**). The decay rate of the P/E state in PostHC (k_off_=3.7 s^-1^**; Table S1**) is much higher than the subunit rotation rate, which is in fact negligible, suggesting that also for deacylated tRNA the hybrid state formation and subunit rotation are not tightly coupled.

To relate subunit rotation to the presence of RF3–GTP on the ribosome, we measured the dissociation rates of RF3 from PreHC and PostHC via the smFRET signal between RF3–Cy5 and L11-Cy3. Labeling of RF3 did not change its catalytic properties (**Figure 2-Figure Supplement 1A,D**). The mean FRET efficiency is approximately 0.7 in both complexes. The interaction of RF3 with PreHC or PostHC is transient, with similar dissociation rates, 5.4 and 5.9 s^-1^, respectively (**Figure 3C,F; Table S1**). We note that these rates are much higher than the rate of GTP hydrolysis by RF3, about 0.5 s^-1^ (Peske et al., 2014; Shi and Joseph, 2016; Zavialov et al., 2001), which implies that the observed rapid RF3 dissociation is independent of GTP hydrolysis. RF3 release is not directly coupled to subunit rotation, as in the presence of RF3 the majority of PostHC does not show fluctuations between the N and R states (**Figure 3D,F, Table 1**), and the rate of R to N transitions of the PreHC is lower than RF3 dissociation (**Figure 3A,C, Table 1**).

### Interplay between RF1 and RF3

Next, we studied the interplay between RF1 and RF3 during termination. To monitor the interactions with PreHC, we again used the RF1(GAQ) mutant, which ensures that peptidyl-tRNA in PreHC is not hydrolyzed. While RF1(GAQ) alone stabilizes the N state (**Figure 4A**, blue line; **Figure 1B**), and RF3 alone induces transitions to the R state (**Figure 3A**), in the presence of RF1(GAQ) and RF3 together the N and R states are almost equally populated (**Figure 4A**). About 60% of the ribosomes show rapid reversible N to R transitions indicating that RF3 can promote subunit rotation even when RF1 is present. RF1(GAQ) binds to a state with a FRET efficiency of 0.7 and its dissociation rate is <0.3 s^-1^ (**Figure 4B,D; Table S1**). RF3 can associate with and dissociate from PreHC–RF1 (**Figure 4C**). The FRET efficiency for the RF3–L11 pair is closer to 0.5, changed from 0.7, which is observed in the absence of RF1 (**Figure 3C**). Thus, the orientation of RF3 in the complex differs from that formed in the absence of RF1, whereas the position of RF1 appears unchanged, at least with respect to L11. The rate of RF3 dissociation is 1.3 s^-1^, which is higher than the dissociation rate of RF1(GAQ), but about 5-fold slower than that of RF3 in the absence of RF1(GAQ) (**Table S1**), indicating that RF1 stabilizes the binding of RF3 to PreHC. Dwell time distributions of the N and R states in the presence of RF1(GAQ) and RF3 are biphasic, suggesting the presence of two populations of each complex. The majority of ribosomes display rapid transitions (>70%, k_N→R_ = 5.9 s^-1^, k_R→N_ = 2.9 s^-1^; **Figure 4D; Table S1**) that are faster than RF1 or RF3 dissociation, suggesting that ribosome fluctuations occur while both factors are bound to the ribosome. Additional low rotation rates (k_N→R_ = 1.3 s^-1^, k_R→N_ = 0.8 s^-1^) may represent spontaneous subunit rotation after factor dissociation (**Table S1**).

**Figure 4.**
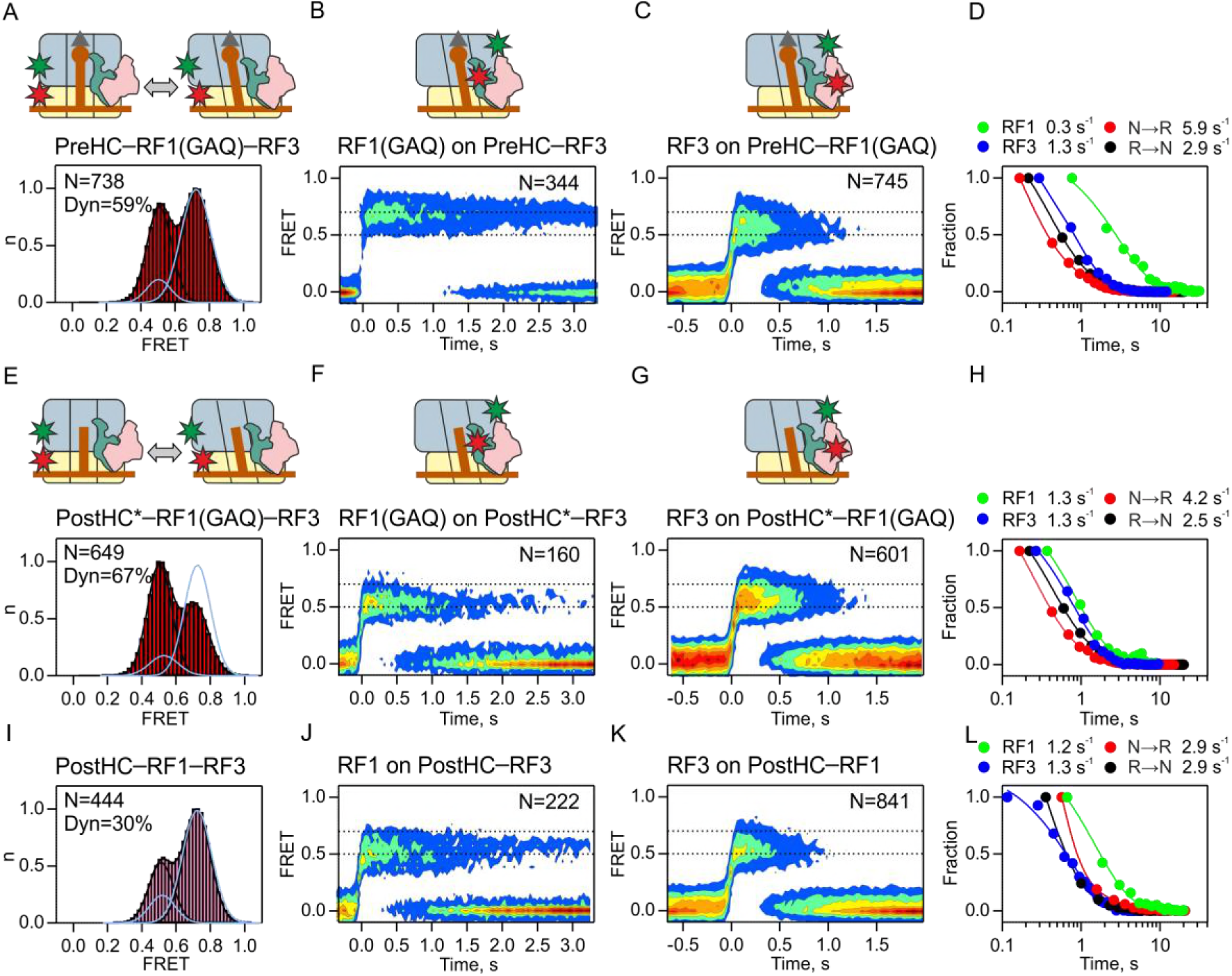
Interplay between RF1 and RF3. (A-D) Interactions with PreHC. (E-H) Interactions with PostHC* obtained by puromycin treatment. (I-L) Interactions with PostHC which is formed *in situ* using RF1. (A,E,I) Subunit rotation measured at saturating factor concentrations (1 μM each). N is the number of traces in the histogram. Blue line represents the distribution on PreHC-RF1 in the absence of RF3. Dyn indicates the percentage of dynamic complexes. (B,F,J) Contour plot representing the residence time of RF1 on the ribosomes monitored by FRET between RF1-Cy5 or RF1(GAQ)-Cy5 (10 nM) and PreHC-Cy3 or PostHC-Cy3 in the presence of excess RF3 (1 μM). Time courses were synchronized to the beginning of the FRET signal. N is the number of traces entering the contour plot.(C,G,K) Contour plot representing the residence time of RF3 on the ribosome monitored by FRET between RF3–Cy5 (10 nM) and PreHC-Cy3 or PostHC-Cy3 in the presence of excess RF1 or RF1(GAQ) (1 μM). (D,H,L) Comparison of the rates of RF1 and RF3 dissociation and subunit rotation. *See also Table S1*.

Next, we monitored subunit rotation in PostHC. For better comparison with the results obtained with PreHC and RF1(GAQ), we first prepared PostHC by puromycin treatment and studied its interactions with RF1(GAQ) and RF3–GTP (**Figure 4E**). The majority (70%) of complexes undergo N to R transitions and the equilibrium is shifted towards the R state. The mean FRET efficiency between RF1(GAQ)-Cy5 and Cy3-labeled L11 is 0.5 (**Figure 4F**), changed from 0.7 on PreHC-RF3 (**Figure 4B**) or on PreHC and PostHC in the absence of RF3 (**Figure 2A-C**). This suggests that peptide release induces a rearrangement of the complex which alters the position of RF1 relative to L11, whereas the position of RF3 is determined by the presence of RF1 (**Figure 3F and 4G**). The dissociation rates are 1.3 s^-1^ for both RF1(GAQ) and RF3 (**Figure 4F-H; Table S1**). A small portion (8%) of complexes that release RF1(GAQ) slowly (k_off_ <0.1 s^-1^) is likely due to incomplete peptide hydrolysis by puromycin. The rotation rates (k_N→R_ = 4.2 s^-1^, k_R→N_ = 2.5 s^-1^) are somewhat higher than RF1 and RF3 dissociation rates, but the most prominent effect of peptide release is the acceleration of RF1 dissociation from <0.3 to 1.3 s^-1^ (**Figure 4D,H; Table S1**).

A similar dynamic behavior is observed in the presence of wild-type RF1 and RF3 bound to PostHC: the R state is enriched compared to the PostHC with RF1 alone and the complexes show reversible N to R transitions (**Figure 4I**). The rates of RF1 dissociation are k1 = 1.2 s^-1^ (72%) and k_2_ = 0.3 s^-1^ and RF3 dissociation is 1.3 s^-1^, respectively, and the FRET efficiency is 0.5 for both RF1 and RF3 (**Figure 4J-L; Table S1**). The subunit rotation rate is 2.9 s^-1^ in either direction. Thus, RF1 and RF3 can reside simultaneously on the PostHC in an arrangement where the position of RF1 and RF3 relative to L11 is shifted compared to the complex with only a single factor. The ribosomes undergo frequent transitions between the N and R states, but these fluctuations are not directly coupled to the dissociation of the factors.

To understand whether RF3 can dissociate from the N state when RF1 is bound, we used apidaecin 137 (Api), an antibiotic that enters the polypeptide exit tunnel after peptide release, interacts with RF1 and blocks RF1 dissociation from the PostHC (Florin et al., 2017). The PostHC–RF1–Api–RF3 complex is stalled in the N state (**Figure 5A**), whereas in the absence of RF1 Api does not alter the N and R state distribution induced by RF3 (**Figure 5B**). In the PostHC–RF1–Api–RF3 complex RF1 is stably bound in the state with FRET 0.7 relative to L11 (**Figure 5C**). RF3 is bound in the state with FRET 0.5 to L11 and dissociates with the rate of 1.2 s^-1^ (**Figure 5D**).

**Figure 5.**
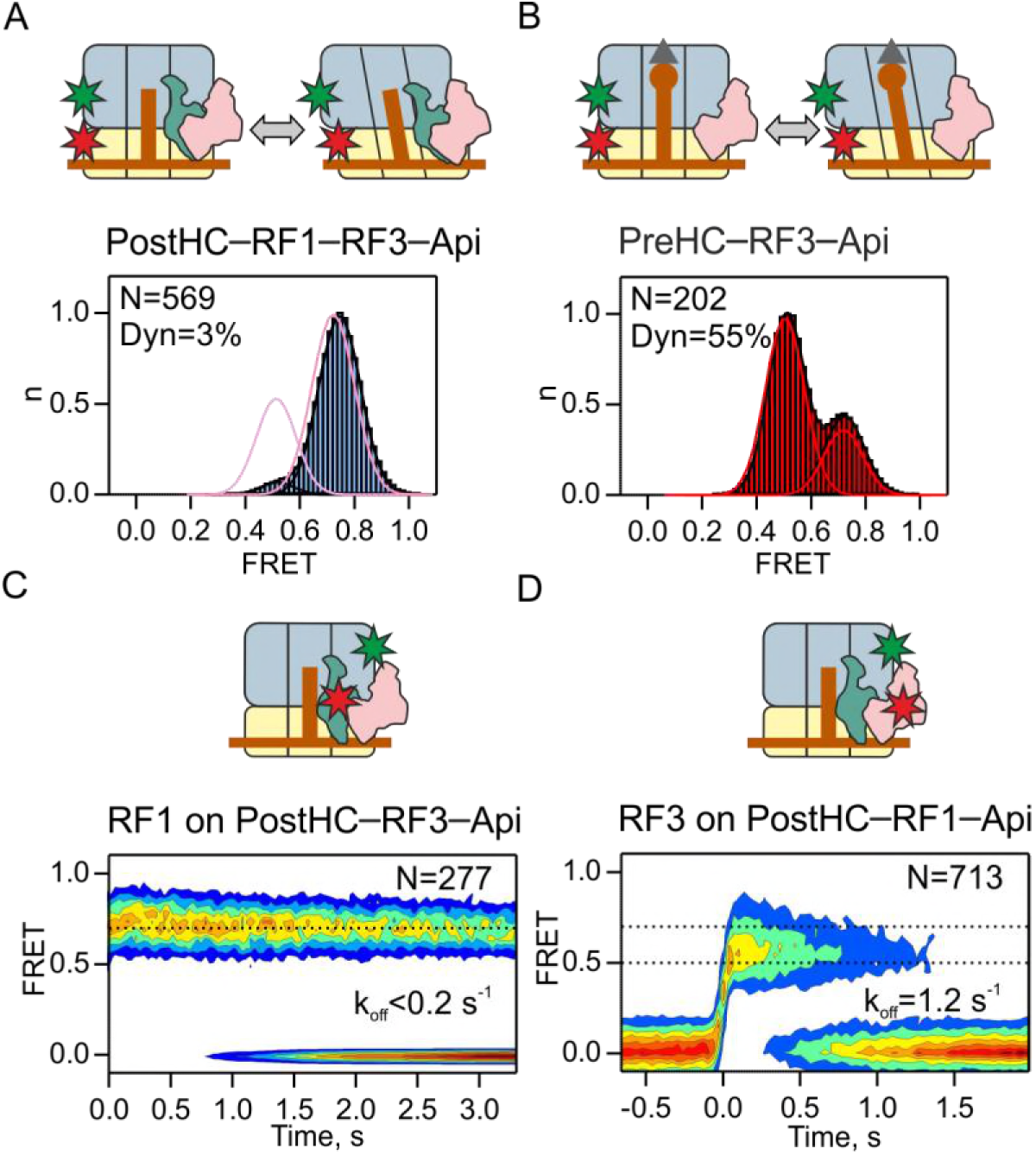
Dissociation of RF3 from the ribosomes in static N state. (A,B) Subunit rotation measured with or without RF1 (1 μM), with saturating RF3 concentrations (1 μM) and Api137 (Api, 1 μM). Pink line in (A) or red line in (B) represents the distribution of states in the absence of Api. N is a number of traces used in the histogram. Dyn indicates the percentage of dynamic complexes. (C) Contour plot representing the residence time of RF1-Cy5 on PostHC-Cy3 in the presence of saturating RF3 (1 μM) and Api (1 μM). (D) Contour plot representing the residence time of RF3–Cy5 on PostHC-Cy3 in the presence of saturating RF1 (1 μM) and Api (1 μM). *See also Table S1*

### The role of GTP binding and hydrolysis

By analogy with other GTPases, GTP hydrolysis by RF3 is expected to regulate the dissociation of RF3 from the ribosome. In contrast to all other GTPases, RF3 was suggested to bind to the PostHC–RF1 complex in the GDP-bound form; the ribosome-induced rapid release of GDP should stabilize RF3 binding, while the subsequent GTP binding induces the conformational change of the ribosome and release of RF1 (Sternberg et al., 2009; Zavialov et al., 2002). We first tested these notions using a biochemical turnover peptidyl-tRNA hydrolysis assay (**Figure 6A,B**). When both RF1 and RF3 are substoichiometric to PreHC, such that 10 cycles of RF1 and RF3 turnover are required to convert all PreHC to PostHC, GTP hydrolysis is essential (**Figure 6A**). In excess of RF3, when only RF1 has to turnover, efficient peptide release is observed with wt RF3 in the presence of GTP, GTPγS or GDPNP (**Figure 6B**). Also RF3(H92A)–GTP, a RF3 mutant deficient in GTP hydrolysis, induces efficient RF1 recycling, contrary to previous reports (Gao et al., 2007), but consistent with a recent kinetic study (Shi and Joseph, 2016). Apo-RF3 has no activity, again consistent with previous reports (Shi and Joseph, 2016; Zavialov et al., 2002). The low activity in the presence of GDP is most likely due to a minor contamination with GTP. Thus, GTP hydrolysis is not required for RF1 recycling when active GTP-bound RF3 is in excess, but is necessary to ensure recycling of RF3, while the apo- and GDP-forms of RF3 appear inactive.

**Figure 6.**
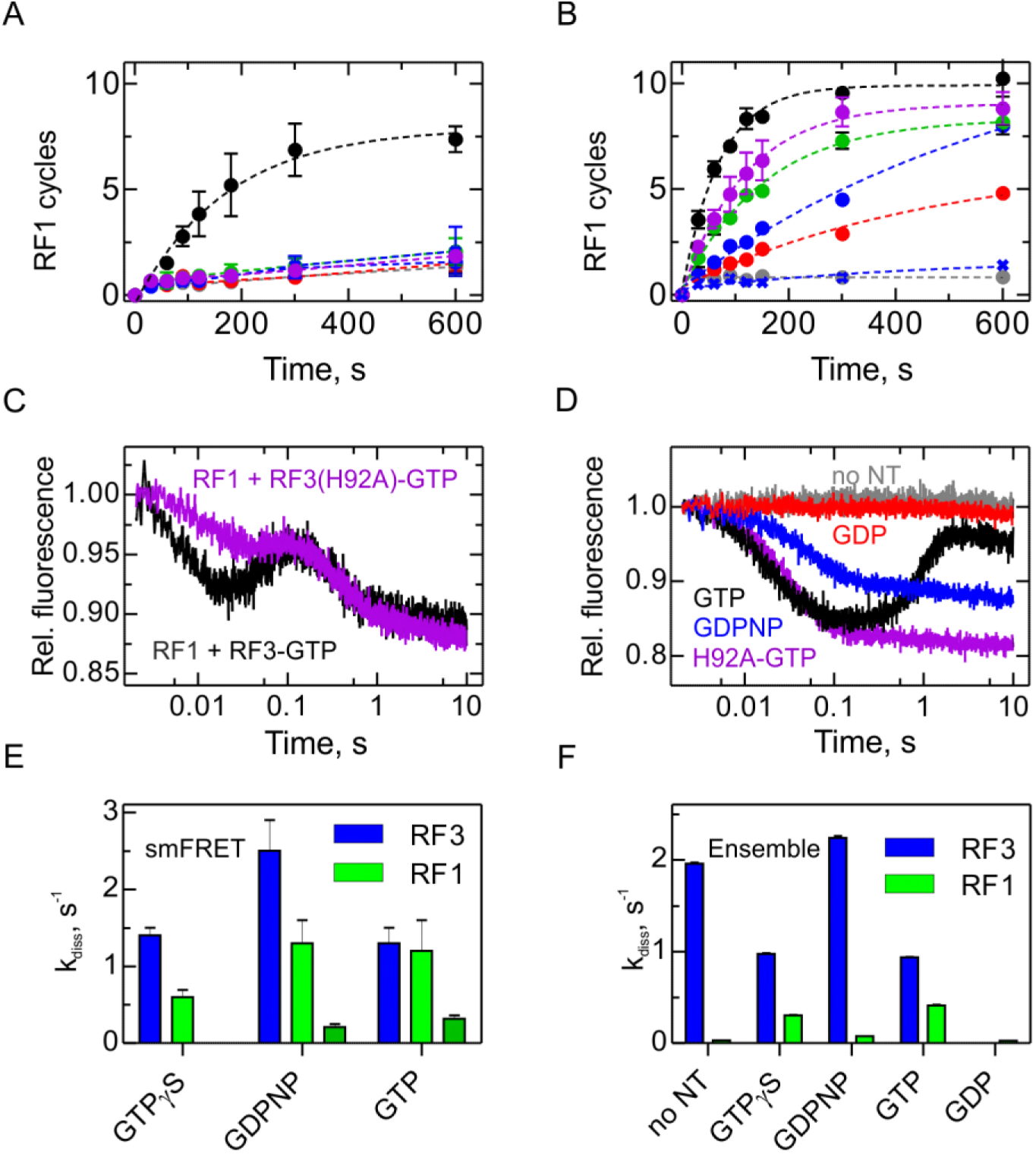
The role of GTP hydrolysis for RF1 and RF3 recycling. (A,B) Effect of different nucleotides on peptidyl-tRNA hydrolysis (GTP, black circles; GTPγS, green circles; GDPNP, blue circles; GDP, red circles; no nucleotide, grey circles) or in the presence of RF3(H92A) and GTP (purple circles). Control experiments are in the absence of RF3 (blue crosses). Error bars represent the range of two technical replicates. (A) Peptide hydrolysis was performed by incubating PreHC (100 nM) with RF3 (10 nM) and the respective nucleotides (1 mM); reactions were started with the addition of RF1 (10 nM). (B) Same as in (A), but at 1 μM RF3. (C,D) Kinetics of subunits rotation induced by release factors. PreHC (50 nM) with fluorescence-labeled S6 and L9 (Sharma et al., 2016) was rapidly mixed with (C) RF1 (1 μM) and either wt RF3 or RF3(H92A) (1 μM) in the presence of GTP (1 mM) or (D) wt RF3 (1 μM) or with RF3(H92A) (1 μM) in the presence of different nucleotides (1 mM) in the absence of RF1. (E,F) Dissociation rates of RF1 (green bars) and RF3 (blue bars) in the presence of different nucleotides measured by smFRET (E) or ensemble kinetics (F). In cases where the dissociation of RF1 is bi-phasic, the second dissociation rate is shown (dark green). Error bars represent the standard error of the exponential fit determined in three independent data sets. *See also Figure 6-Figure Supplement 1, Figure 6-Figure Supplement 2, Figure 6-Figure Supplement 3, Figure 6-Figure Supplement 4 and Table S1*

Next, we sought to understand the role of GTP binding for the conformational dynamics of the ribosome. In the apo- or GDP-bound form, RF3 does not induce the shift from N to R state that is observed with RF3–GTP, and the ribosomes remain static (**Figure 6-Figure Supplement 1A-D**); this observation holds for both PreHC–RF3 and PostHC–RF1–RF3. By analogy, smFRET experiments monitoring the position of the L1 stalk show that addition of RF3–GDP or apo–RF3 does not change the ribosome conformation (Sternberg et al., 2009). For a more direct observation of RF3 binding to the ribosome, we then used the RF3–Cy5 and L11-Cy3 FRET pair and studied RF3 binding to PostHC–RF1, but we did not find any FRET events characteristic of RF3 binding in the presence of GDP or with apo-RF3 (data not shown). It is known that RF3–GDP or apo-RF3 can bind to PostHC–RF1 in some way, because this interaction accelerates nucleotide exchange in RF3 (Koutmou et al., 2014; Peske et al., 2014; Shi and Joseph, 2016; Zavialov et al., 2001). However, the interaction must be transient and does not engage the factor at its binding site at L11 unless GTP is bound.

We then asked whether GTP hydrolysis by RF3 is required to induce subunit rotation (**Figure 6-Figure Supplement 1E-G**). We used three different ways to block GTP hydrolysis: by utilizing RF3(H92A) and by replacing GTP by a non-hydrolysable analogs, GDPNP, which is often applied in structural studies, or GTPγS, which appears more similar to GTP. At saturating concentrations of RF3(H92A)–GTP, the distribution of N and R states in PreHC in the absence of RF1 is identical to that with the wt RF3–GTP (**Figure 6-Figure Supplement 1E** and **Figure 3A**). In the presence of RF3–GDPNP, the R state is less favored, whereas with RF3–GTPγS the state distribution is more similar to that with GTP (**Figure 6-Figure Supplement 1F,G**). This may occur if different non-hydrolysable analogs induce slightly different conformations of the complex independent of GTP hydrolysis or if the affinity of the RF3–GDPNP or RF3–GTPγS to the ribosome is reduced. The conformational dynamics of PostHC–RF1 follows the same tendency: the state distribution is similar with GTP and RF3–GTPγS and shifts slightly towards the N state with GDPNP (**Figure 6-Figure Supplement 2A,E**). In all cases the fraction of dynamic ribosomes is somewhat reduced. Thus, the authentic GTP-form of RF3, but not GTP hydrolysis, is required to induce the shift in the distribution of the N and R states observed at steady-state conditions (**Figure 4I**).

Because the smFRET experiments alone did not explain the role of GTP hydrolysis, we monitored the kinetics of ribosome rotation by rapid kinetics using the same labeling positions on the proteins S6 and L9, but a different yet well-characterized FRET dye pair which is better suited for the stopped-flow setup (Ermolenko and Noller, 2011; Sharma et al., 2016). Upon mixing of PreHC with RF1 and RF3, subunits rapidly rotate from N towards R state (**Figure 6C**) due to peptide release and factor dissociation. The time courses are multiphasic, suggesting stepwise rearrangement in the complexes. The slower accumulation of the intermediates between 0.01 and 0.1 s is likely due to slightly reduced affinity of RF3(H92A) to the ribosome compared to the wild-type RF3. The predominant late kinetic phase and the final FRET level reflecting the fraction of R to N state are independent of GTP hydrolysis, in agreement with the smFRET steady-state data (**Figure 6-Figure Supplement 2**).

**Figure 7.**
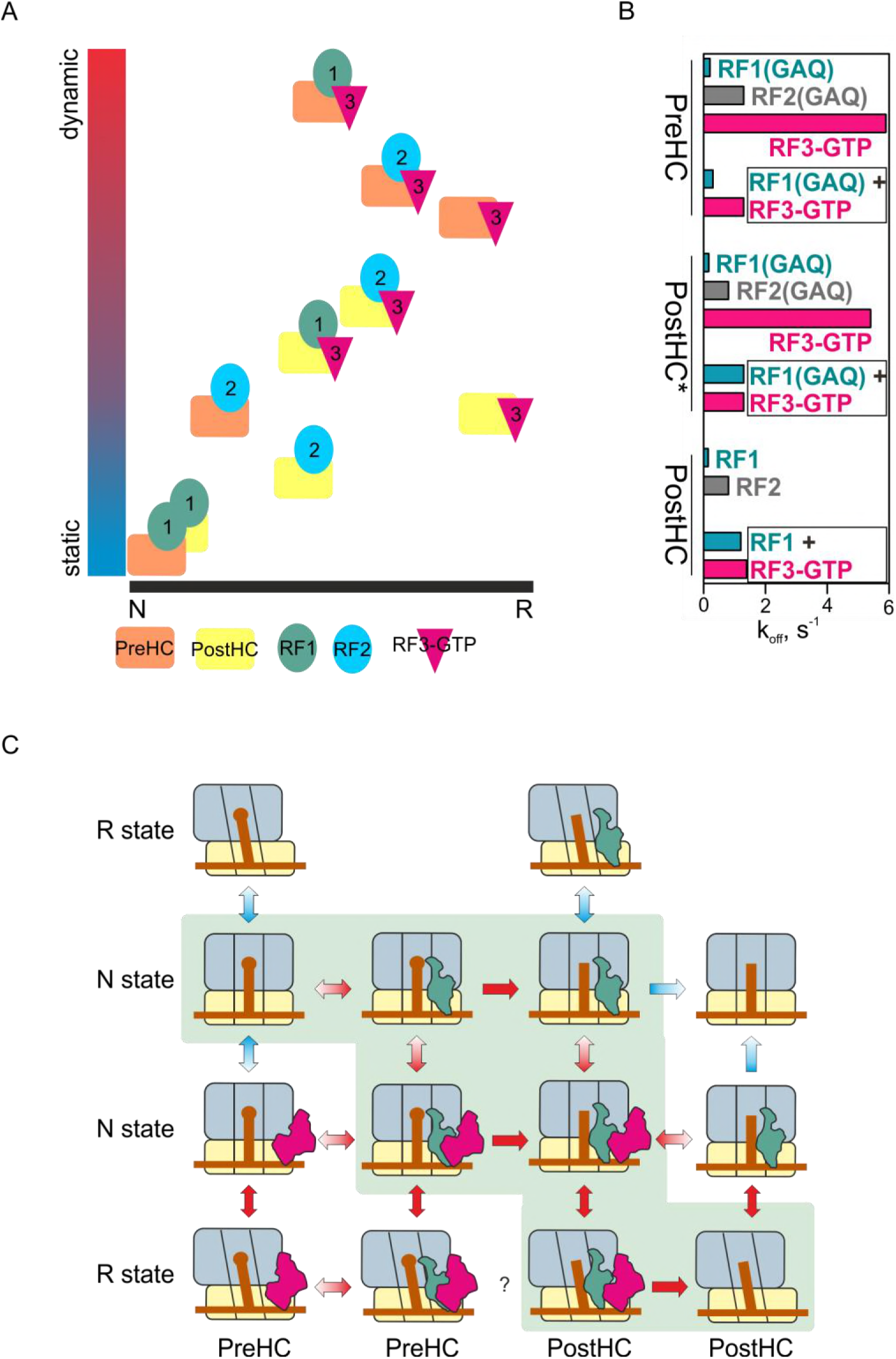
The mechanism of translation termination. (A) Ribosome dynamics in the presence of RF1, RF2, and RF3. Ribosome fluctuations are color-coded from static (blue) to dynamic (red) and correlated to the fraction of N and R state in the respective complex. (B) Summary of the dissociation rate constants of RF1, RF2 and RF3 from different type of complexes. Bars representing the dissociation of RF1 are colored teal, RF2 gray, RF3 magenta. (C) The landscape of ribosome conformations with RF1 and RF3. The ribosome states (N and R, PreHC and PostHC) are indicated. Red arrows indicate rapid reaction, blue arrows static or slowly exchanging states, with the preferential direction indicated by color gradient; black arrows indicate peptidyl-tRNA hydrolysis, with the asterisk indicating peptide release from the R state, the efficiency of which is not clear. *See also Table S1*

We next asked whether the dissociation rates of RF1 and RF3 are affected by GTP hydrolysis. We addressed the question by two approaches, by smFRET using the RF-L11 FRET pair and by ensemble kinetics stopped-flow technique using other FRET and fluorescence reporters than in the smFRET experiments (**Figure 6-Figure Supplement 3**). Depending on the GTP analog, RF1 adopts a conformation with FRET 0.7 or 0.5 relative to L11 (**Figure 6-Figure Supplement 2B,F,I**). The dwell time distributions for RF1 show heterogeneous kinetics with the rates of the rapid phase around 0.6 s^-1^ to 1.3 s^-1^, respectively, and a slower phase of about 0.2 to 0.3 s^-1^ (**Figure 6E; Table S1**); the latter slow phase is monitored by ensemble kinetics (**Figure 6F**). Surprisingly, the observed high rate of RF3 dissociation is either not affected (1.4 s^-1^ with GTPγS) or is even higher (2.5 s^-1^ with GDPNP) than with GTP (**Figure 6E**). The rates of RF3 dissociation are very similar in ensemble and smFRET experiments and are independent of GTP hydrolysis (**Figure 6E,F**). Again, the apo and GDP-forms of RF3 were inactive in stimulating RF1 dissociation. These data suggest that GTP hydrolysis by RF3 does not regulate RF3 dissociation from the PostHC–RF1 complex. Rather, non-hydrolysable analogs have an indirect effect on how RF3 modulates conformational preferences of RF1 (**Figure 6-Figure Supplement 2B,F**).

Given that the experiments performed in the presence of RF1 did not reveal a significant effect of GTP hydrolysis, we performed the ensemble ribosome rotation experiments with PreHC in the absence of RF1 (**Figure 6D**). PreHC in the absence of RF3 is predominantly in the N state and should return into the same state after RF3 dissociation, whereas RF3 binding induced a strong accumulation of the R state (**Figure 6D**). With GTP, a fraction the complex remained in the R state for a relatively long period and then rotated back slowly. A similar behavior is seen with PostHC–RF3; in the latter case the amplitude is lower, because the portion of complexes in the N state is smaller than in PreHC (data not shown). This result is unexpected, because the dissociation of RF3–GTP as measured by smFRET is rapid (~6 s^-1^; **Figure 3C,F**) and does not explain the long persistence of the stalled R state. In the absence of GTP hydrolysis, the N state is not recovered, suggesting that the ribosomes are blocked in the R state, presumably bound to RF3 (**Figure 6D**) (Gao et al., 2007; Jin et al., 2011; Koutmou et al., 2014; Zhou et al., 2012). One possibility is that a fraction of complexes remains stalled in the R state with RF3 and resolving these complexes requires GTP hydrolysis which is slow (0.5 s^-1^; (Peske et al., 2014; Shi and Joseph, 2016; Zavialov et al., 2001; Zavialov et al., 2002)). In contrast, dissociation from the N state (or from the complexes that contain RF1) does not require GTP hydrolysis and is rapid (**Figure 5D, 6E,F,** and **Figure 6-Figure Supplement 2C,G**). Thus, based on the results of experiments with RF3(H92A), GTP hydrolysis by RF3 is required to accelerate the factor dissociation from those ribosomes that do not contain RF1 and are trapped in the R state. Depletion of the RF3 pool by the R-state ribosomes that do not carry RF1 accounts for the observed inhibition of RF1 turnover in the termination assay (**Figure 6A**).

This model predicts that the RF3 dissociation from the R-state ribosomes in the absence of RF1 should be reduced compared to the experiments with RF1 (**Figure 6E,F**). This was in fact observed in both smFRET and ensemble experiments (**Figure 6-Figure Supplement 4**). However, we note that the number of smFRET trajectories was much smaller compared to the same type of experiments with RF3–GTP, and the rapid phase of RF3 dissociation was not found (**Table S1**). It is possible that the affinity of the RF3 with non-hydrolysable analogs is reduced due to very rapid dissociation of RF3 from the N state, which makes these complexes too transient to be observed and explains why only the slow dissociation rates are seen. We note, however, that GTP analogs may induce conformations that are dissimilar from the authentic GTP-bound state, which calls for some caution in interpreting data obtained with non-hydrolysable GTP analogs.

## Discussion

Our experiments show how release factors navigate through the landscape of possible ribosome conformations during translation termination. The range of conformations sampled by the ribosome during termination encompasses the static N state, dynamic states fluctuating between N and R with a preference to one or another state, and the static R state (**Figure 7A**). Release factors not only change the distribution between the N and R states, but also alter the fraction of the ribosomes that make transient fluctuations between the states. Binding of RF1 to either PreHC or PostHC favors the static N state; in the PostHC—RF1 complex protein L1 adopts an open conformation and the tRNA appears to be in a classical state (Sternberg et al., 2009), which is coupled to the formation of the N state. The N state of the ribosome–RF1 complex is also captured by structural studies (James et al., 2016; Korostelev et al., 2008; Laurberg et al., 2008; Petry et al., 2005; Weixlbaumer et al., 2008). Surprisingly, we find that PreHC with RF2 is more dynamic, and has higher fraction of the R states than the complex with RF1. Furthermore, RF2 can dissociate equally well from the PreHC and PostHC and is less dependent on the action of RF3 than RF1 (**Figure 2-Figure Supplement 3E**, **Figure 7B**). The ability of RF2 to function is only regulated by the ratio between the rate of peptide release and factor dissociation, e.g., if the rate of peptidyl-tRNA hydrolysis is about 10 s^-1^ at cellular pH (Indrisiunaite et al., 2015; Kuhlenkoetter et al., 2011) and the rate of RF2 dissociation is ~1 s^-1^ (this paper), the factor can achieve efficient peptide release before dissociating. Thus, RF1 and RF2 – albeit fulfilling a similar function during termination – differ in their ability to affect ribosome dynamics.

Binding of RF3–GTP to PreHC or PostHC induces the rotation toward the R state (**Figure 7A**). The PreHC–RF3 complex is dynamic, whereas the PostHC–RF3 is stabilized in a static R state, consistent with the previous smFRET work (Sternberg et al., 2009) and structural studies (Gao et al., 2007; Jin et al., 2011; Zhou et al., 2012). After peptide release RF1–RF3 or RF2–RF3 together shift the distribution of ribosome conformations towards the middle of the dynamic spectrum (**Figure 7A**). The rates of ribosome fluctuations are in the range of 2–6 s^-1^, somewhat faster than in the absence of the factors, 0.5–2.6 s^-1^ (**Table S1**).

One open question is what drives the dissociation of RF1 and RF3 from the ribosome (**Figure 7B**). Dissociation of RF1 from the static N state is slow; the acceleration by RF3 correlates with increased ribosome dynamics and frequent transitions from N to R state. However, dynamic transitions alone are not sufficient to induce RF1 dissociation from the ribosome, because peptide release is crucial to allow RF1 to dissociate rapidly. Peptide release correlates with a change in the orientation of RF1 with respect to L11. On the other hand, peptide release alone is also not sufficient, as the dissociation rate of RF1 from PostHC is slow in the absence of RF3. Thus, RF1 dissociation is promoted by the concerted action of RF3 (which stimulates subunit rotation and may directly displace RF1 from its binding site) and by peptide release (which allows for a conformational adjustment of RF1).

RF3 dissociation is independent of peptide release, but is affected by the presence of RF1 or RF2, which stabilize RF3 binding to the ribosome and change conformation/position of RF3 relative to L11. In the presence of RF1, RF3 efficiently dissociates from the N state even in the absence of GTP hydrolysis (this paper and (Shi and Joseph, 2016)). The order of RF1 and RF3 dissociation appears random, because the rates of factor release are quite similar and the exact sequence depends on experimental conditions (this paper; (Koutmou et al., 2014; Shi and Joseph, 2016)). In those cases where RF1 happens to dissociate before RF3 has left the ribosome, GTP hydrolysis completes RF3 recycling. In summary, subunit rotation, peptide release, conformational changes of the factors, and GTP hydrolysis together drive dissociation of RF1 and RF3. However, kinetically these movements are not directly coupled, i.e. the dissociation rates of the factors and the rates of subunit rotation are independent of each other, but are individually defined by the dynamic properties of the complex.

This work provides an unexpected view on the role of nucleotide exchange and GTP hydrolysis by RF3. Although RF3-GDP or apo-RF3 can bind to the ribosome carrying RF1/RF2 (Peske et al., 2014; Zavialov et al., 2001), this interaction does not result in the recruitment of the factor at its binding site at the vicinity of L11. *In vitro* in the absence of GTP, apo-RF3 can form a relatively stable complex with PostHC–RF1 (Pallesen et al., 2013; Shi and Joseph, 2016), but this binding does not alter the dynamics of subunit rotation and does not accelerate RF1 dissociation (this paper and (Sternberg et al., 2009)). Rather, the GTP-bound form of RF3 is required to stimulate ribosome dynamics and RF1 dissociation from PostHC (**Figure 6A,B**) (Zavialov et al., 2001; Zavialov et al., 2002). At the high concentrations of GTP in the cell, apo-RF3 will be immediately converted into the functionally active GTP form; thus, the apo-RF3–ribosome complex is a transient intermediate. Furthermore, given the moderate difference in the affinities of RF3 for GTP and GDP at cellular concentrations, a large fraction of RF3 is in the GTP form. The present experiments, most of which are performed in the presence of the GTP-regeneration system and thus do not allow for accumulation of the GDP- or apo-form of RF3, show efficient factor binding, peptide release and factor recycling. We thus have no indication for an active role of nucleotide exchange or for the essential role of the GDP- or apo-form of RF3 in termination at cellular conditions and we consider the respective models unlikely.

Unexpectedly, our data suggest that GTP hydrolysis is primarily required to release RF3 from a portion of the ribosomes that are arrested in the R state in the absence of RF1/RF2, whereas other ribosomes can release RF3 even in the absence of GTP hydrolysis. This finding explains why PostHC, with its higher propensity to be in the R state than the PreHC, is more efficient in stimulating GTP hydrolysis by RF3 (Zavialov et al., 2002). In this respect, RF3 appears to be an unusual GTPase that differs from other translational GTPases, such as EF-G, EF-Tu and IF2, where GTP hydrolysis is coupled to key steps on the reaction pathway of the factors and is required on all ribosome complexes. Rather, the internal clock of the RF3 GTPase (Peske et al., 2014) acts as a rescue mechanism to release prematurely recruited RF3 to the complexes that do not contain RF1. This scenario is realistic at the concentrations of factors in the cell where RF3 is much more abundant than RF1 (Schmidt et al., 2016). On the other hand, RF3 is reminiscent of other GTPases in that GTP hydrolysis controls a conformational rearrangement of the ribosome. In the physiological setting, the R state that forms after peptide release and dissociation of RF1/RF2 is favored for the subsequent recruitment of RRF and EF-G, resulting in ribosome recycling (Prabhakar et al., 2017).

These results lead to the following model of translation termination for RF1 (**Figure 7C**). RF1 and RF3-GTP can bind to any conformation of PreHC. Among all possible reaction routes, two appear most likely, either via RF1 binding, then peptide release, then RF3-GTP recruitment, or through simultaneous binding of RF1 and RF3–GTP to PreHC followed by peptide release. This central complex PostHC–RF1–RF3–GTP fluctuates between N and R states and allows for rapid dissociation of both RF1 and RF3. Multiple ribosome conformations, ribosome dynamics and the lack of strong coupling between the rates of subunit rotation and the dissociation of RF1 and RF3 seem characteristic features of RF1-dependent termination. Thus, translation termination is a stochastic process that utilizes loosely coupled motions of its players to complete protein synthesis and release the newly synthesized nascent chain towards its cellular destination.

## AUTHOR CONTRIBUTIONS

All authors designed the experiments and discussed results. S.A. performed all smFRET experiments; S.A. and T.S. analyzed the data; C.M., H.S., P.K. and W.H. prepared materials and performed biochemical and ensemble kinetics experiments; I.W. carried out mass spectrometry analysis; S.A., C.M., T.S., F.P., and M.V.R. interpreted the results. S.A. and M.V.R. wrote the paper with contributions of all authors.

## ACKNOWLEDGMENTS

We thank Marija Liutkute for preparation of RF2-Cy5 and Olaf Geintzer, Franziska Hummel, Sandra Kappler, Christina Kothe, Anna Pfeifer, Theresia Uhlendorf, Tanja Wiles, and Michael Zimmermann for expert technical assistance. We thank Dr. H. Urlaub and the bioanalytical mass spectrometry facility for assistance with the analysis of RF1/RF2 methylation. The work was supported by the grants of the Deutsche Forschungsgemeinschaft (SFB860 for M.V.R.).

## Materials and Methods

### CONTACT FOR REAGENT AND RESOURCE SHARING

*Further information and requests for resources and reagents should be directed to and will be fulfilled by the Lead Contact, Marina V. Rodnina*.

### METHOD DETAILS

#### Buffers

All smFRET experiments were performed in imaging buffer (50 mM Tris-HCl pH 7.5, 70 mM NH_4_Cl, 30 mM KCl, 15 mM MgCl_2_, 1 mM spermidine, 8 mM putrescine, 2.5 mM protocatechuic acid, 50 nM protocatechuate-3,4-dioxygenase (from *Pseudomonas*), 1 mM Trolox (6-hydroxy-2,5,7,8-tetramethylchromane-2-carboxylic acid), and 1 mM methylviologen. Rapid kinetics and peptide hydrolysis experiments were performed in TAKM_7_ buffer (50 mM Tris-HCl pH 7.5, 70 mM NH_4_Cl, 30 mM KCl, 7 mM MgCl_2_).

#### Labeled ribosomes and release factors

The preparation and functional characterization of ribosomes labeled with Cy3 at protein L11 and double-labeled at S6-Cy5 and L9-Cy3 was carried out as described (Adio et al., 2015; Sharma et al., 2016). RF2 construct was cloned from the *E. coli* K12 strain and contains the natural T246A replacement (Wilson et al., 2000). Catalytically impaired RF1(G234A) (RF1(GAQ)) and RF2(G251A) (RF2(GAQ)), and the respective single-cysteine variant RF1(S167C) (Sternberg et al., 2009; Wilson et al., 2000), RF2(C273) and RF3(L233C) were generated by Quickchange mutagenesis according to the standard protocol. Native cysteines were replaced by serine or alanine based on the sequence conservation analysis performed using the Consurf database. RF1 and RF2 were purified and *in vitro* methylated as described (Kuhlenkoetter et al., 2011). RF3 was purified by affinity chromatography on a Ni-IDA column (Macherey-Nagel) followed by ion exchange chromatography on a HiTrapQ column (Peske et al., 2014).

Prior to labeling, methylated RF1 and RF2 were incubated for 30 min with a 10-fold molar excess of TCEP (Sigma) at room temperature (RT). Fluorescein (Sigma), QSY9 (Thermo Scientific) and Cy5 (GE Healthcare) maleimides were dissolved in DMSO and added to the proteins (5- to 10-fold molar excess). Labeling was performed for 2 hour at RT and quenched by addition of a 10-fold molar excess of 2-mercaptoethanol over dye. Excess dye was removed by gel filtration on a PD-10 column (GE Healthcare).

#### mRNA

All mRNAs used in the smFRET experiments are labeled with biotin at the 5'end and were purchased from IBA (Göttingen, Germany). The following sequences were used:

mMetStop

5'-Biotin-CAACCUAAAACUUACACACCCGGC<u>AAGGAGG</u>UAAAUA**AUG**<u>UAA</u>ACG AUU-3'

mMetPheStop

5'-Biotin-CAACCUAAAACUUACACACCCGGC<u>AAGGAGG</u>UAAAUA**AUG**UUU<u>UAA</u>ACGAUU-3'

mMetLysStop

5'-Biotin-CAACCUAAAACUUACACACCCGGC<u>AAGGAGG</u>UAAAUA**AUG**AAG<u>UAA</u>ACGAUU-3'

mMetValStop

5'-Biotin-CAACCUAAAACUUACACACCCGGC<u>AAGGAGG</u>UAAAUA**AUG**GUU<u>UAA</u>ACGAUU-3'

For the peptide hydrolysis and rapid kinetics experiments, ribosome complexes were assembled on the synthetic model mRNA, 5'-GGCAAGGAGGUAAAUA**AUG**<u>UAA</u>ACGAUU-3' (IBA) with a start codon followed by a stop codon.

#### Sample preparation for smFRET TIRF experiments

Initiation complex formation was carried out by incubating ribosomes (100 nM) with a three-fold excess of IF1, 2 and 3, fMet-tRNA^fMet^, mRNA and 1 mM GTP in TAKM_7_ for 30 min at 37°C. In case of the mRNA coding for fMetStop, the initiation complex was used as PreHC. To generate PreHC on other mRNAs, an equal volume of ternary complex was added containing EF-Tu (1 μM) incubated with GTP (1 mM), phosphoenolpyruvate (3 mM) and pyruvate kinase (0.1 mg/ml) in TAKM_7_ for 15 min at 37°C, followed by addition of Phe-tRNA^Phe^, Lys-tRNA^Lys^ or Val-tRNA^Val^ (500 nM). Addition of EF-G (100 nM) and GTP (1 mM) induced tRNA translocation to form PreHC that contains peptidyl tRNA in the P site and displays the UAA stop codon in the A site.

#### TIRF experiments

Complexes were diluted to 1 nM with smFRET buffer (50 mM Tris-HCl, 70 mM NH_4_Cl, 30 mM KCl, 15 mM MgCl_2_, 1 mM spermidine and 8 mM putrescine). Biotin/PEG functionalized cover slips were incubated for 5 min at room temperature with the same buffer containing additionally BSA (10 mg/ml) and neutravidin (1 μM) (Thermo Scientific). Excess neutravidin was removed by washing the cover slip with buffer containing BSA (1 mg/ml). Ribosome complexes were applied to the surface and immobilized through the mRNA-biotin:neutravidin interaction. Images were recorded at a rate of 30 frames/s after exchanging the buffer with imaging buffer at room temperature (22°C) (Adio et al., 2015).

To monitor subunit rotation of L9/S6-labeled ribosomes in the presence of release factors at steady-state, imaging buffer was supplemented with RF1, RF2 and/or RF3 (1 μM each). In experiments with RF1(GAQ) or RF2(GAQ) the observation time was limited to <10 min in order to minimize peptide hydrolysis due to residual factor activity. In experiments monitoring subunit rotation by RF3 in the GTP form or in complex with non-hydrolysable GTP analogs, imaging buffer was additionally supplemented with the energy recycling system (1 mM GTP or 1 mM GDPNP or 1 mM GTPγS, 3 mM phosphoenolpyruvate and 0.1 mg/ml pyruvate kinase). FRET signals reporting on the time course of subunit rotation during termination were obtained by injecting RF1 or RF2 (100 nM) in imaging buffer to immobilized PreHC or PostHC.

To measure FRET signals reporting on the residence time of labeled release factors on PreHC or PostHC labeled at protein L11 with Cy3, the complexes were immobilized on the cover slip. Movies were recorded upon addition of Cy5-labeled RF1, RF2 or RF3 to a final concentration of 10 nM in imaging buffer. To study the residence time of Cy5-labeled RF1 or RF1(GAQ) on ribosomes in the presence of RF3, imaging buffer was supplemented with unlabeled RF3 (1 μM), GTP (1 mM), phosphoenolpyruvate (3 mM) and pyruvate kinase (0.1 mg/ml). To study the residence time of Cy5-labeled RF3 in the presence of RF1, imaging buffer was supplemented with unlabeled RF1 (1 μM), GTP (1 mM), phosphoenolpyruvate (3 mM) and pyruvate kinase (0.1 mg/ml).

To monitor FRET signals reporting on the conformation of the P-site tRNA PreHC or PostHC labeled on protein L1(C202-Cy5) and on fMet-tRNA^fMet^(thioU8-Cy3) or tRNA^fMet^(U8-Cy3) were immobilized on the coverslip. Movies were recorded upon addition of imaging buffer or imaging buffer containing RF2(GAQ) (1 μM) or 1RF3 (1 μM). In experiments monitoring subunit rotation by RF3 imaging buffer was additionally supplemented with the energy recycling system (1 mM GTP or 1 mM GDPNP or 1 mM GTPγS, 3 mM phosphoenolpyruvate and 0.1 mg/ml pyruvate kinase) (Sternberg et al., 2009).

#### Data Analysis

Fluorescence time courses for donor (Cy3) and acceptor (Cy5) were extracted using custom made Matlab (MathWorks) software (Adio et al., 2015). A semi-automated algorithm (Matlab) was used to select anti-correlated fluorescence traces exhibiting characteristic single fluorophore fluorescence intensities. Acquired time traces were selected from the data set by choosing only traces that contained single photobleaching steps for Cy3 and Cy5. The bleed-through of the Cy5 signal into the Cy3 channel was corrected using an experimentally determined coefficient (~0.13 in our experimental system). All trajectories were smoothed over three data points. FRET efficiency was defined as the ratio of the measured emission intensities, Cy5/(Cy3 + Cy5). FRET-histograms were fitted to Gaussian distributions using Matlab code.

The vbFRET software package (http://vbfret.sourceforge.net/) (Bronson et al., 2009) was used for hidden Markov model (HMM) analysis of the FRET data. FRET probability density plots are two-dimensional contour maps generated from time-resolved FRET traces. For the experiments measuring subunit rotation of PostHC upon binding of RF1 in real time, FRET traces were synchronized at the transition entering the stable N state. For the experiments measuring subunit rotation of PreHC upon binding of RF2 in real time, FRET traces were synchronized to the first N to R transition. In experiments measuring the residence time of labeled release factors FRET traces are synchronized to the beginning of the FRET event reporting on the binding of the factor to the ribosome. Dwell times of different FRET states of fluctuating traces were calculated from idealized traces. Dwell time histograms were fitted to either one- or two-exponential function. Rates (k) were calculated by taking the inverse of dwell times. Rate constants and errors were determined from three independent data sets as described (Fei et al., 2011; Wasserman et al., 2016). Statistics is provided in Table S1.

#### Peptide hydrolysis assay

PreHC was prepared as described (Peske et al., 2014) and purified through sucrose cushion centrifugation. After centrifugation, ribosome pellets were resuspended in TAKM_7_, frozen in liquid nitrogen and stored at -80°C. The extent of initiation was better than 95% as determined by nitrocellulose filtration and radioactive counting. [^3^H]PreHC (100 nM) was incubated with RF3 at the indicated concentration and nucleotide (1 mM) for 15 min at 37°C. Pyruvate kinase (0.1 mg/ml) and phosphoenol pyruvate (3 mM) were added in all experiments performed in the presence of GTP. Time courses were started by addition of RF1 or RF2 (10 nM). Samples were quenched with a solution containing TCA (10%) and ethanol (50%). After centrifugation (30 min, 16,000 g) the amount of released f[^3^H]Met in the supernatant was quantified by radioactive counting.

#### Rapid kinetics

Rapid kinetic experiments were performed on an SX-20MV stopped-flow apparatus (Applied Photophysics, Leatherhead, UK), by rapidly mixing equal volumes (60 μl) of reactants at 37°C. Dissociation of RF1 from the ribosome was monitored in chase experiments using complexes containing fluoresceine-labeled tRNA^fMet^ (Flu) in PostHC-Flu prepared by incubation of PreHC-Flu (50 nM) with RF1-QSY9 (50 nM). The complex was rapidly mixed with RF3 (1 μM) and unlabeled RF1 (500 nM). RF3–Flu (100 nM) dissociation was monitored by mixing PostHC (100 nM), RF1 (1 μM) and RF3–Flu (100 nM) with an excess of unlabeled RF3 (1 μM). Fluorescein was excited at 470 nm and monitored after passing a KV500 filter (Schott, Mainz, Germany). Subunit rotation experiments were carried out using double-labeled ribosomes (S6-Alx488/L9-Alx568)(Sharma et al., 2016), by rapidly mixing PreHC (50 nM) with release factors (1 μM). Alx488 was excited at 470 nm and fluorescence of Alx568 was monitored after passing through a KV590 cut-off filter (Schott). Time courses were evaluated by exponential fitting using GraphPad Prism software.

## Supplementary Figures

**Figure 1-Figure Supplement 1.**
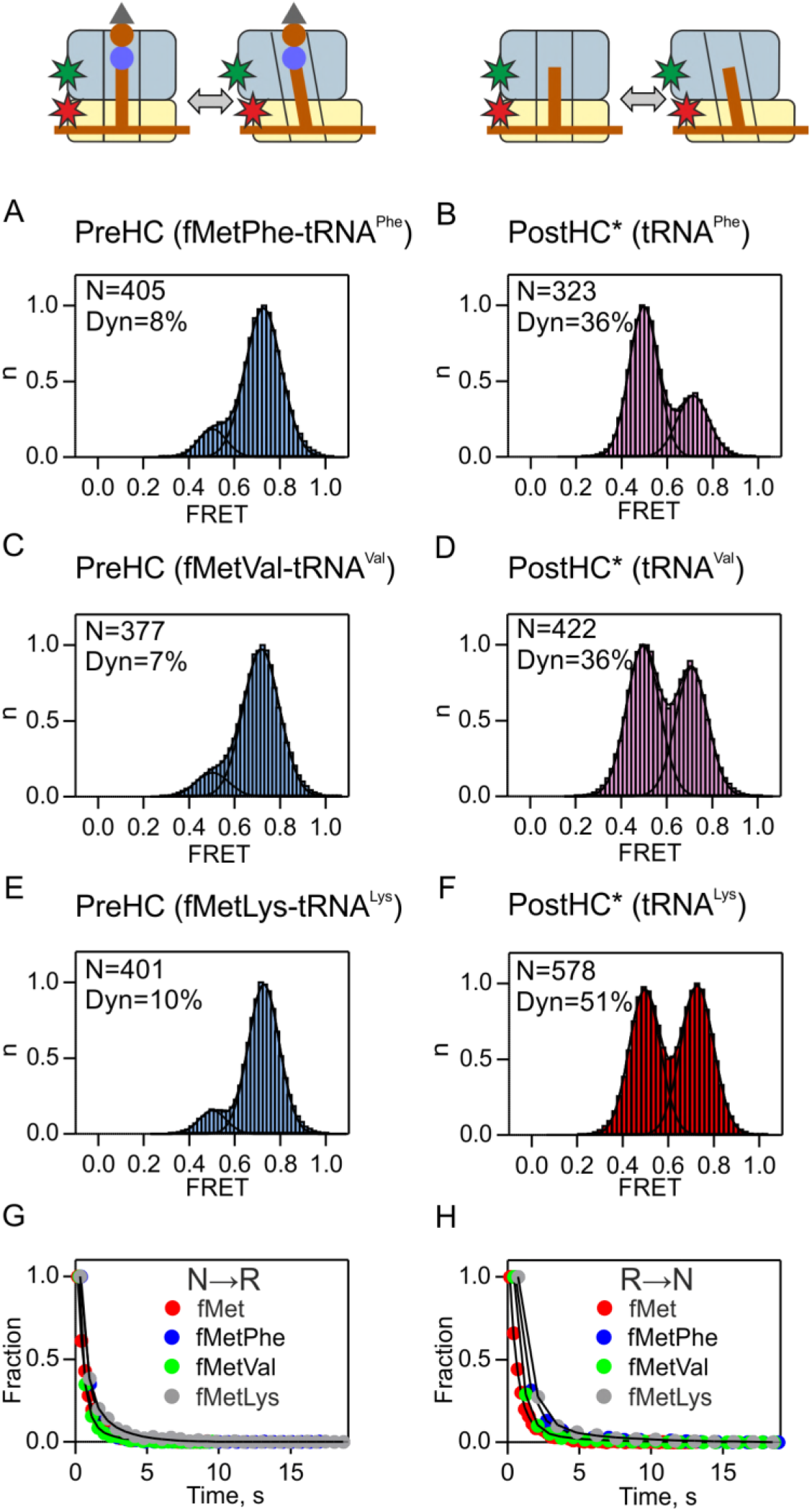
Subunit rotation monitored by FRET between S6-Cy5 and L9-Cy3. PreHC (A,C,E) and PostHC* (B,D,F) with different tRNAs in the P site. PostHC* was generated by addition of puromycin (1 mM) to PreHC. Histograms show the normalized distribution of FRET states. N is the number of traces entering the histogram. Dyn indicates the percentage of dynamic complexes. (G,H) Dwell time distributions of the N to R and R to N transitions. For the rates see the Table S1.

**Figure 1-Figure Supplement 2.**
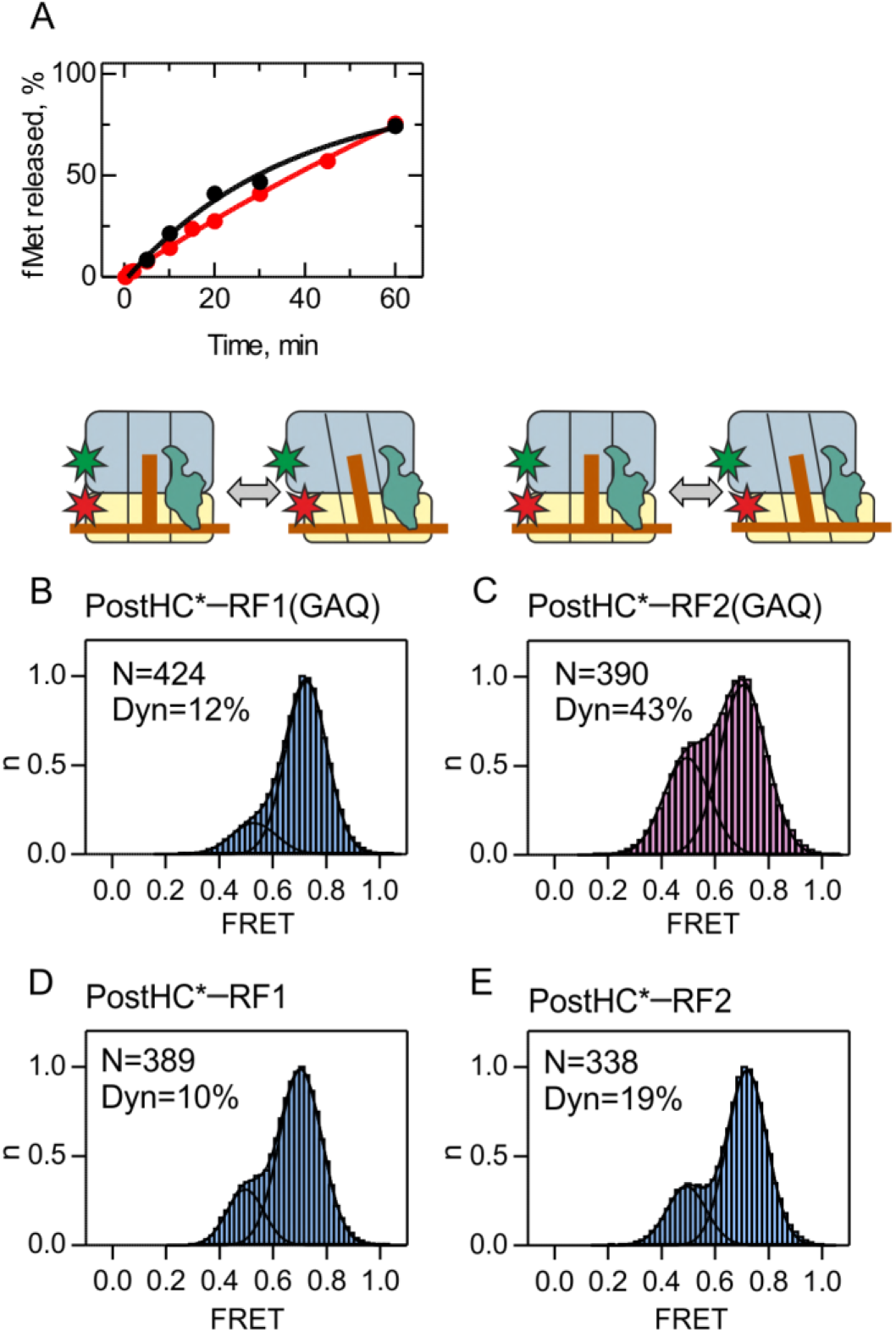
Peptide hydrolysis by RF1/RF2(GAQ) mutants and subunit rotation of Post HC* monitored using FRET between S6-Cy5 and L9-Cy3. (A) Time courses of fMet-tRNA^fMet^ hydrolysis by RF1(GAQ) (black) and RF2(GAQ) (red) at 23°C in TAKM_7_ buffer. All smFRET experiments were completed in less than 10 min after mixing. (B,D) In the presence of RF1 or RF1(GAQ) (1 μM). (C,E) In the presence of RF2 or RF2(GAQ) (1 μM). *PostHC was generated using puromycin

**Figure 1-Figure Supplement 3.**
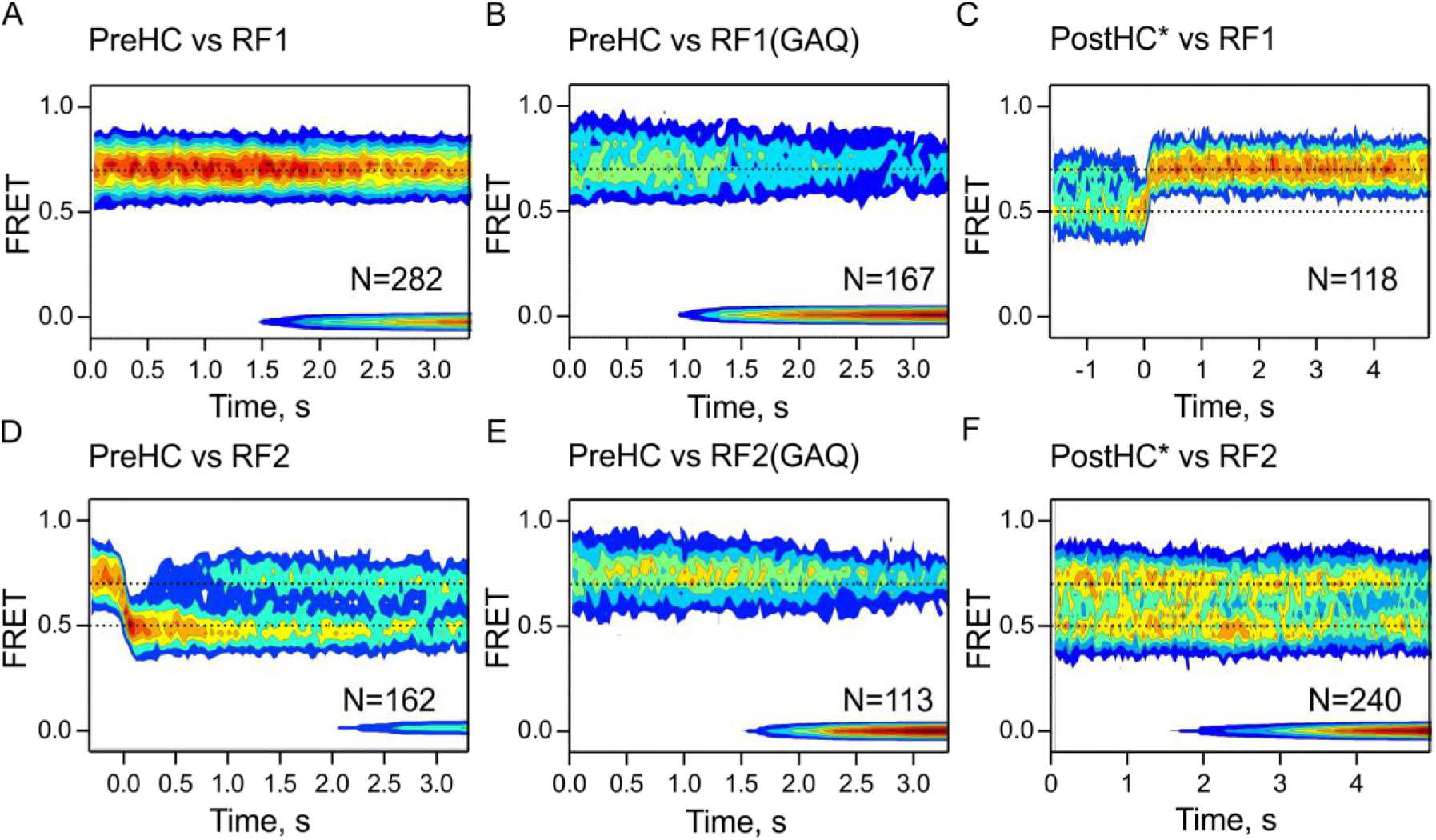
Time-resolved subunit rotation. The experiments were performed in the presence of low concentrations (100 nM) of RF1 or RF1(GAQ) (A-C) or RF2 or RF2(GAQ) (D-F). All time traces are combined in contour plots. In (C) traces are synchronized to the last R to N transition. In (D) traces are synchronized to the first N to R transition. Dashed lines indicate the mean FRET values for the N state (0.7) and R state (0.5). N is the number of traces entering the contour plot.

**Figure 2-Figure Supplement 1.**
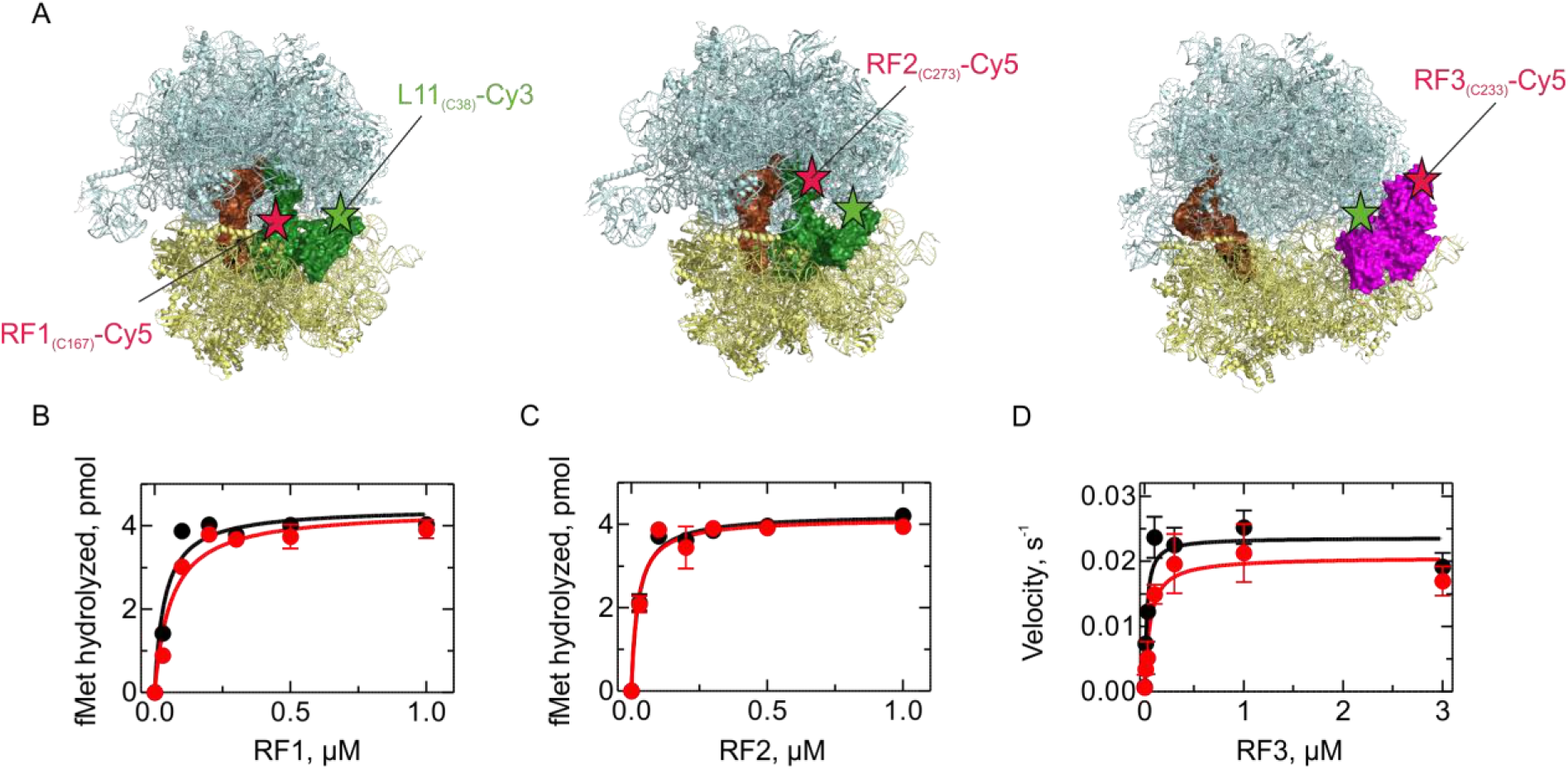
Activity of the fluorescence-labeled RFs. (A) Labeling positions in RF1, RF2, and RF3 (indicated as red stars) and L11 (green stars) Structural models were prepared using PDB entries 4V7P (Korostelev et al., 2010) (RF1), 5CZP (Pierson et al., 2016) (RF2) and 4V89 (Zhou et al., 2012) (RF3). (B,C) Activity of RF1-Cy5 (B) and RF2-Cy5 (C) (red) compared to the respective wild-type factor (black) as determined by peptide hydrolysis. PreHC (30 nM) was incubated with the indicated concentrations of RF for 10 s. (D) Activity of RF3–Cy5 (red) compared to the wild-type factor (black) as determined by the ability to recycle RF1. PreHC (100 nM) was incubated with RF1 (10 nM) and increasing concentrations of RF3. The rate of peptide hydrolysis was determined by linear fitting of the time courses at initial velocity conditions.

**Figure 2-Figure Supplement 2.**
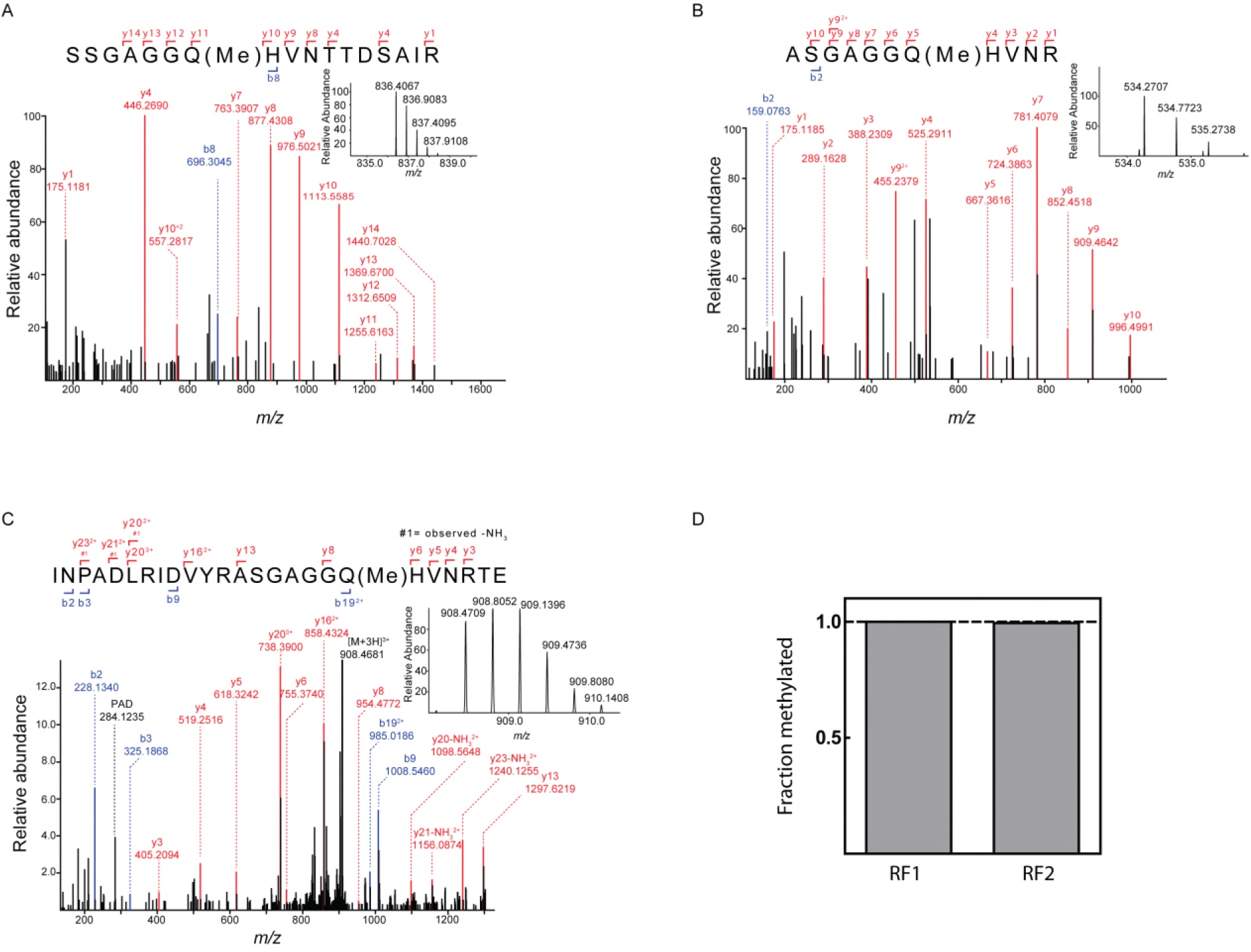
Quantification of release factor methylation by mass spectrometry. (A) MS/MS spectrum of methylated peptide derived from RF1-Cy5 by proteolysis with trypsine. Inset: MS spectrum *(m/z* = 836.4059) of the intact methylated peptide (z=2). (B) MS/MS spectrum of the methylated peptide derived from RF2-Cy5 by proteolysis with trypsine. Inset: MS spectrum *(m/z* = 534.2707) of the intact methylated peptide (z=2). The exceptional hydrophilicity of the peptide hampered a reliable relative quantification. (C) MS/MS spectrum of methylated peptide derived from RF2-Cy5 by proteolysis with GluC. Inset: MS spectrum (m/z=908.4706) of the intact methylated peptide (z=3). (D) Relative quantification of the methylation efficiency. Methylated and unmethylated peptide was quantified by MS1 full scan filtering and the fraction of methylated peptide was quantified assuming similar ionization properties of both peptides. Mean values were calculated from 3 technical replicates with error bars (±0.001) representing standard deviation (too small to be seen).

**Figure 2-Figure Supplement 3.**
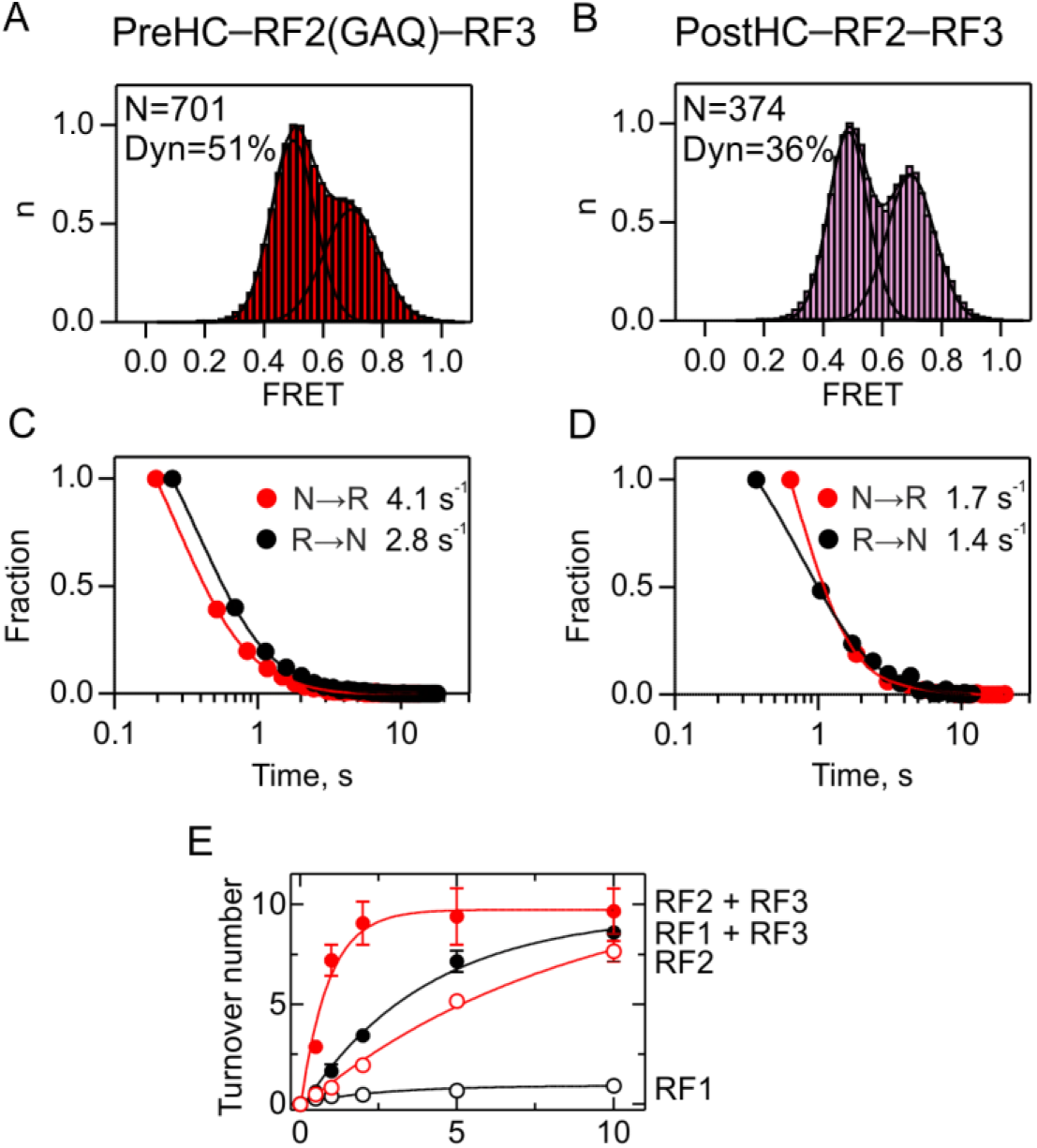
Interplay between RF2 and RF3. (A,B) Subunit rotation with RF2 and RF3–GTP for PreHC (A,C) and PostHC (B,D). (C,D) Dwell time distributions of the N and R states for PreHC (A,C) and PostHC (B,D). N is the number of traces entering the histogram. Dyn indicates the percentage of dynamic complexes. (E) RF3–mediated recycling of RF1 and RF2. Peptide hydrolysis by RF1 (black symbols) and RF2 (red symbols) was monitored at turnover conditions in the absence (open symbols) or in the presence (closed symbols) of RF3. PreHC (100 nM) was incubated with RF1 or RF2 (10 nM) or with RF1 or RF2 (10 nM), RF3 (100 nM) and GTP (1 mM). Error bars represent the standard error of three independent replicates.

**Figure 3-Figure Supplement 1.**
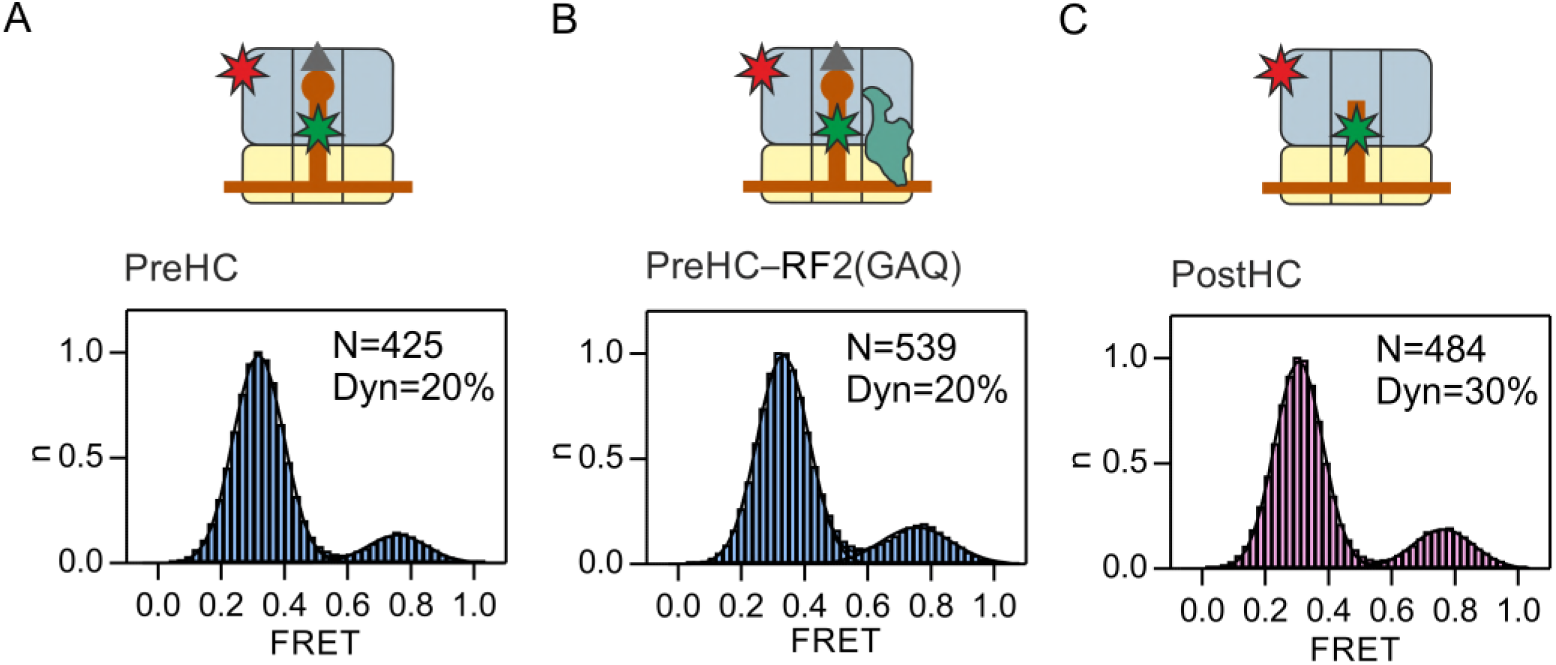
Open and closed conformations of the ribosome measured with an L1-tRNA FRET pair. FRET distribution measured with the fMet-tRNA^fMet^(Cy3)-L1(Cy5) FRET pair in PreHC in the absence of the release factors (A), in the presence of RF2(GAQ) (1 μM) (B), or in PostHC in the absence of the factors (C).

**Figure 6-Figure Supplement 1.**
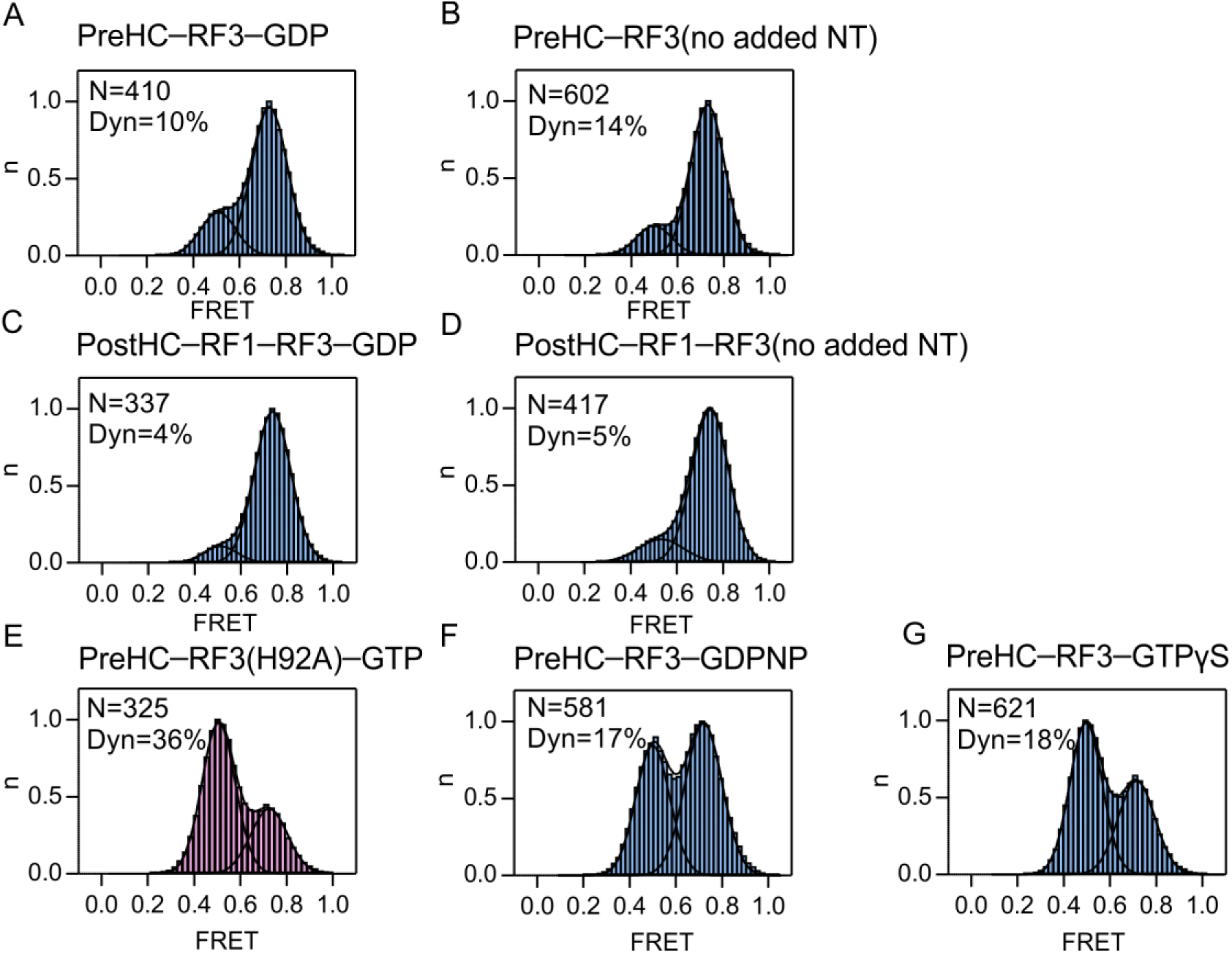
Effect of the nucleotide bound to RF3 on subunit rotation. (A,C) RF3 in the presence of GDP in the absence (A) or presence (C) of RF1. (B,D) Apo-RF3 in the absence of added nucleotide in the absence (B) or presence (D) of RF1. (E) RF3(H92A) and GTP. (F) RF3 with GDPNP. (G) RF3 with GTPγS. In (E,F,G) the GTP regeneration system (phosphoenolpyruvate and pyruvate kinase) was added. The concentration of RF3 is 1 μM and of the nucleotide 1 mM. N is the number of traces entering the histogram. % Dyn indicates the percentage of dynamic complexes.

**Figure 6-Figure Supplement 2.**
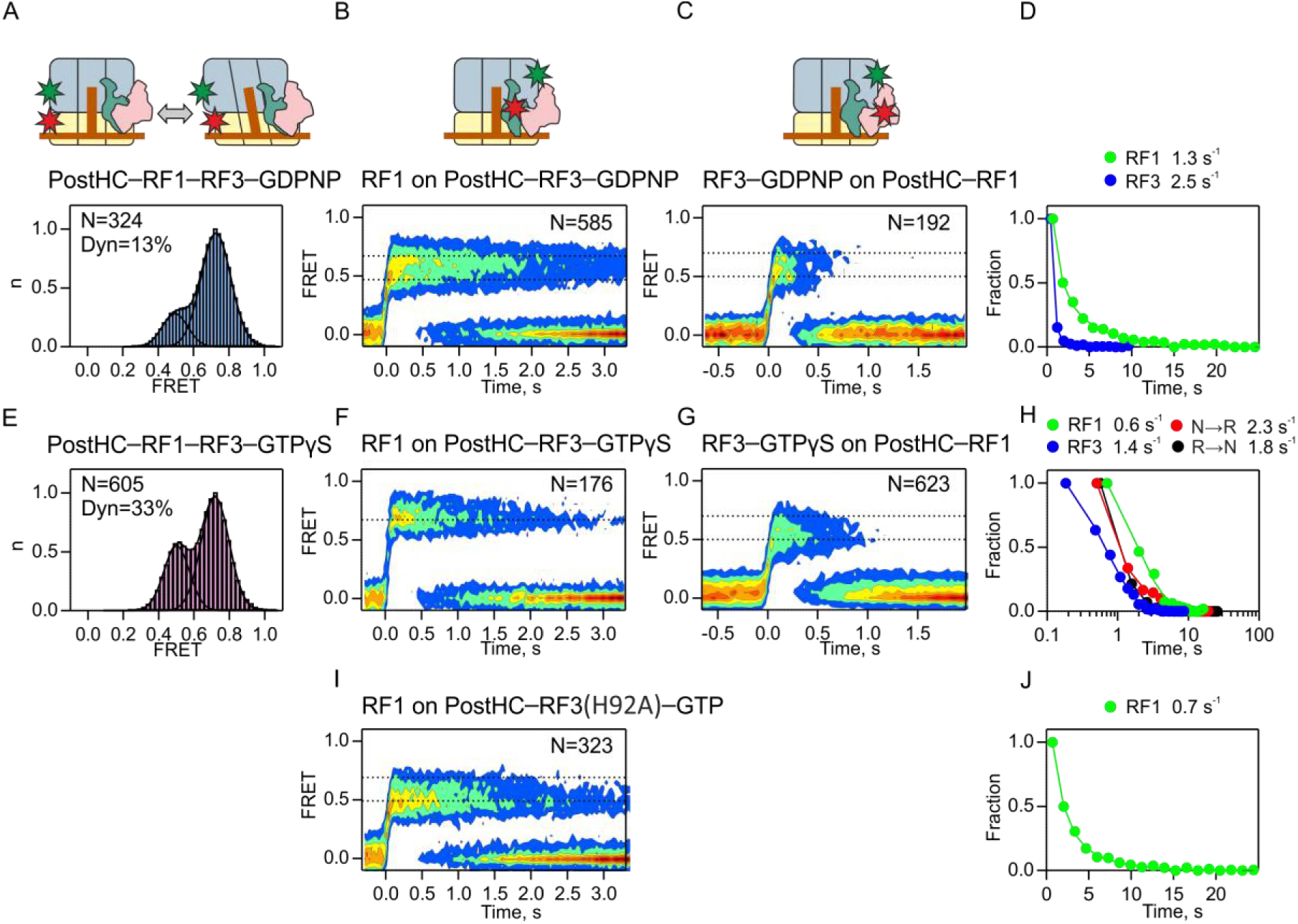
Interplay between RF1 and RF3 on PostHC in the absence of GTP hydrolysis. (A-D) In the presence of GDPNP. (E-H) In the presence of GTPγS. (I,J) With RF3(H92A) and GTP. (A,E) Distribution of N and R states at steady-state conditions with saturating factor concentrations (1 μM each). N is a number of traces used in the histogram. Dyn indicates the percentage of dynamic complexes. (B,F,I) Dissociation of RF1 monitored by FRET between RF1-Cy5 (10 nM) and L11-Cy3 in the presence of excess RF3 (1 μM). Time courses were synchronized to the onset of the FRET event and combined in contour plots. N is the number of traces entering the contour plot. (C,G) Dissociation of RF3 monitored by FRET between RF3–Cy5 (10 nM) and L11-Cy3 in the presence of excess RF1 (1 μM); the experiment with RF3(H92A) was not done owing to the lack of fluorescence-labeled mutant protein. (D,H,J) Dwell time distributions and the rates of RF1 and RF3 dissociation and subunit rotation.

**Figure 6-Figure Supplement 3.**
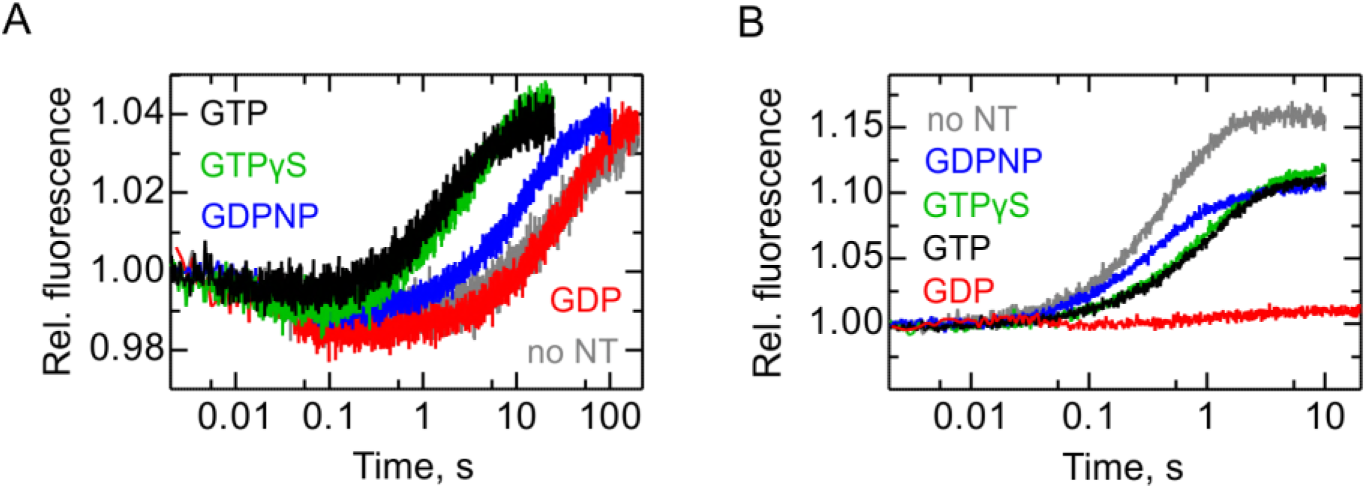
Kinetics of RF1 and RF3 dissociation from the ribosome measured by ensemble kinetics. (A) Time courses of RF1 dissociation measured by stopped flow upon mixing PostHC–RF1 (50 nM) carrying FRET labels on RF1 (RF1-QSY9) and tRNA^fMet^ (fluorescein) with RF3 (1 μM) and unlabeled RF1 (500 nM). The traces shown represent the average of six to eight technical replicates. (B) Time courses of RF3 dissociation upon mixing PostHC–RF1–RF3 (100 nM) carrying fluorescein-labeled RF3 (100 nM) with unlabeled RF3 (1 μM). The traces shown represent the average of five to nine technical replicates.

**Figure 6-Figure Supplement 4.**
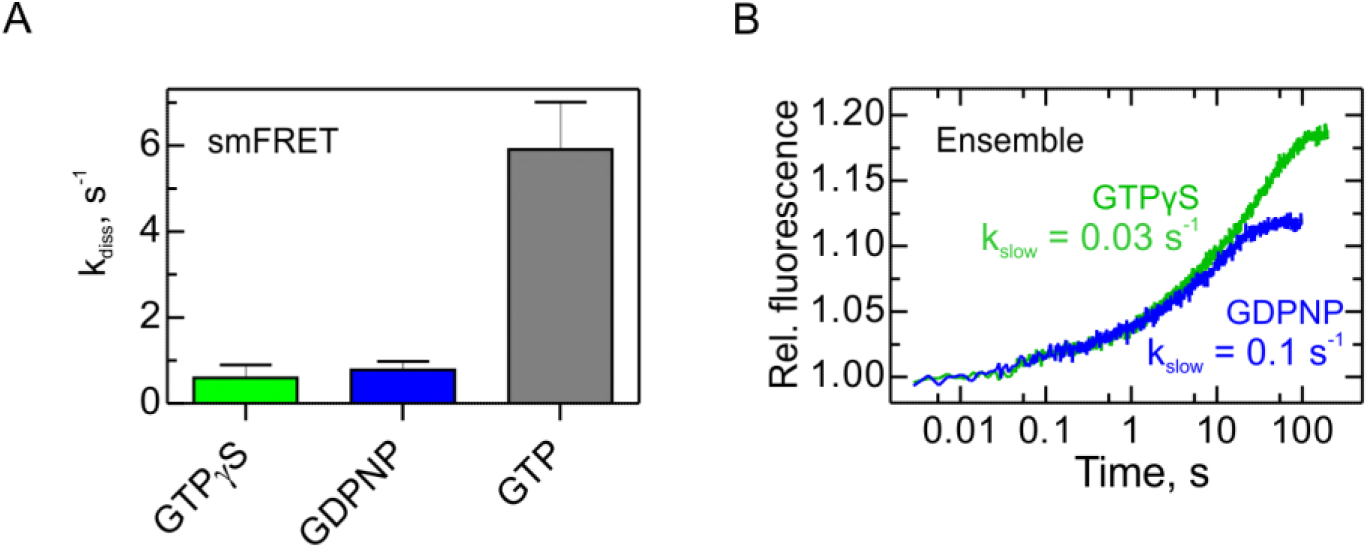
RF3 dissociation from PreHC in the absence of RF1. (A) Dissociation rates of RF3–Cy5 from PreHC-Cy3 derived from smFRET experiments; the experiments were performed as in Figure 3. (B) Time courses of RF3–Flu (100 nM) dissociation fromPreHC-RF3 (100 nM) in the absence of RF1 upon chasing with unlabeled RF3 (1 μM). The traces shown are the average of five to six technical replicates.

**SI Table S1. Related to Figures 1–7.** Quantitative analysis of conformational dynamics during termination. This table is a separate Excel file.

